# GABAergic neurons in the dorsal raphe nucleus regulate social hierarchy in mice

**DOI:** 10.1101/2024.09.17.613408

**Authors:** Lidi Lu, Yanzhu Fan, Shaoxiang Ge, Yue Wu, Zhiyue Wang, Tao Qing, Suxin Shi, Guangzhan Fang

## Abstract

Social hierarchy serves as a fundamental organizational mechanism within most animal societies, exerting significant influence on health, survival, and reproductive success in both humans and animals. However, the neural mechanisms by which the brain regulates dominance hierarchies remain inadequately understood. Considering that GABAergic neurons in the dorsal raphe nucleus (DRN) exert substantial inhibitory control over serotonergic firing, which may be implicated in the acquisition of dominance, we hypothesized that DRN GABAergic neurons may play a pivotal role in regulating social hierarchy. To test this hypothesis, we employed a combination of optogenetics, chemogenetics, fiber photometry recordings, and behavioral assays in mice, to elucidate the functional contributions of these neurons. Our results revealed a biphasic activity pattern of DRN GABAergic neurons, characterized by increased firing during retreats and decreased firing during push-initiation in the tube test. Furthermore, the optogenetic and chemogenetic activation of DRN GABAergic neurons led to an increase in the number of retreats and a reduction in social rank, while inhibition of these neurons produced the opposite effects. These findings elucidate the bidirectional regulatory role of DRN GABAergic neurons in social hierarchies.

## Introduction

The dominance hierarchy servers as a fundamental organizational mechanism within most animal societies^1^. An individual’s position within this hierarchy is often associated with a range of fitness-related outcomes. For example, this social structure influences access to resources and significantly affects health, survival, and reproductive success in both humans^2, 3^ and animals^4, 5^. At the population level, all group members derive benefits from the stability provided by dominance hierarchies^6^, which can mitigate agonistic interactions among all individuals, thereby conserving energy and diminishing the risk of physical injury^7^. Nevertheless, the neural mechanisms through which the brain regulates dominance hierarchies remain inadequately understood.w

The regulation of social hierarchy is a multifaceted process that engages various brain regions, neural circuits, and neurotransmitter systems. Among these brain regions^8–10^, the prefrontal cortex (PFC) is identified as the principal regulatory center for social hierarchy^8^. It integrates information regarding social status from multiple brain areas and influences dominant behaviors through its projections to several downstream regions, including the dorsal raphe nucleus (DRN), basolateral amygdala (BLA), hypothalamus, striatum, and periaqueductal gray matter (PAG)^8, 11–15^. The DRN serves as the primary source of serotonin (5-HT) within the brain^16, 17^, highlighting the crucial role of 5-HT neurons in regulating diverse physiological functions and mental disorders^18–28^. The serotoninergic system is closed linked to social rewards and the expression of social hierarchies across various species, such as crustaceans, reptiles, rodents, and primates^9, 29–38^. For example, the injection of 5-HT into the hemolymph of both lobsters and crayfish diminishes the likelihood of retreat and prolongs the duration of aggressive encounters^18, 39^. Similarly, dominant male vervet monkeys (*Cercopithecus aethiops*) exhibit approximately double the concentration of 5-HT compared to their subordinate counterparts^40^. Furthermore, experimentally enhancing serotonergic activity promotes the acquisition of dominance in monkeys, while reductions in this activity correspondingly diminish dominance^34^. Comparable effects have been reported in humans, where the administration of 5-HT similarly impacts social dominance^41^. These findings collectively highlight the potential involvement of DRN and the serotonergic system in the regulation of social dominance and status.

GABAergic neurons constitute the second most prevalent cell type within the DRN^42, 43^. The amplitude of inhibitory postsynaptic currents elicited by local GABAergic neurons is approximately five times greater than that produced by various long-range afferents, indicating a pronounced inhibition of local GABAergic inputs on 5-HT neurons^25, 27, 42^. Additionally, in vivo stimulation of the mPFC leads to a reduction in the firing rates of DRN 5-HT neurons, suggesting that mPFC axons primarily synapse onto DRN GABAergic neurons, which subsequently inhibit 5-HT neuronal activity^44^. In contrast to the activity of DRN 5-HT neurons, DRN GABAergic neurons are activated by aversive stimuli but are inhibited during reward-seeking behavior, including social interaction^36^. This finding implies that GABAergic neurons may play a complementary role in the modulation of behaviors relative to 5-HT neurons. Supporting this notion, optogenetic activation of DRN GABAergic neurons has been shown to induces aversion and social avoidance in mice^24, 45^. Conversely, optogenetic silencing of DRN GABAergic neurons results in disinhibition of adjacent 5-HT neurons and prevents the acquisition of social avoidance in mice faced with social threat^24^. Thus, it is reasonable to hypothesize that DRN GABAergic neurons are integral to the regulation of social hierarchy.

To examine this hypothesis, we employed optogenetics, chemogenetics, fiber photometry recordings, and behavioral assays to elucidate the role of DRN GABAergic neurons in regulating social hierarchy. Our findings reveal that calcium signals in DRN GABAergic neurons increased during retreats and decreased during the initiation of pushes in the tube test. Notably, the optogenetic and chemogenetic activation of DRN GABAergic neurons led to a decrease in rank within the tube test, whereas their inhibition resulted in an elevation of social hierarchy. This study demonstrates the bidirectional regulatory role of DRN GABAergic neurons in social hierarchy and provides new insights into the neural mechanisms that govern this organizational structure in animal societies.

## Results

### Population activity of DRN GABAergic neurons increased during retreat and decreased during push-initiation in the tube test

To investigate the association between social competition and the activity of GABAergic neurons in the DRN, we employed a standardized behavioral paradigm, specifically the tube test^46^, to simulate a socially competitive environment. Using fiber photometry, we recorded neuronal calcium signals in mice during encountering with their cage mates in the tube test. The behaviors exhibited by the mice in the test can be categorized into push-initiation, push-back, stillness, resistance, and retreat^46^. To capture real-time population activity of DRN GABAergic neurons during competition, we administered an adeno-associated virus (AAV2/9) encoding the fluorescent calcium indicator GCaMP6s with GAD67 promoter (AAV2/9-GAD67-GCaMP6s-3xFLAG) and AAV2/9-GAD67-EGFP-tWPA into the DRN of distinct groups of mice (designated as DRN-GCaMP6s mice and DRN-EGFP mice, respectively; see Fig. 1a-b and Supplementary Fig. 1a). We subsequently implanted a fiber optic cannula into the DRN and secured it to the skull for in vivo fiber photometry recordings during the tube test (Fig. 1c). Compared to DRN-EGFP mice, DRN-GCaMP6s mice exhibited significantly enhanced calcium signals in DRN GABAergic neurons during retreat (Fig. 1d-e), while demonstrating significantly reduced calcium signals during push-initiation (Fig. 1f-g). Collectively, these findings indicate that DRN GABAergic neurons are activated during retreat yet exhibit suppressive activity during push-initiation, suggesting a potential negative correlation between DRN GABAergic neuron activity and social competition. Furthermore, we observed that calcium signals in DRN GABAergic neurons were significantly diminished during effortful behaviors, including push-initiation, push-back, and resistance (Supplementary Fig. 1b-c).

**Figure 1.**
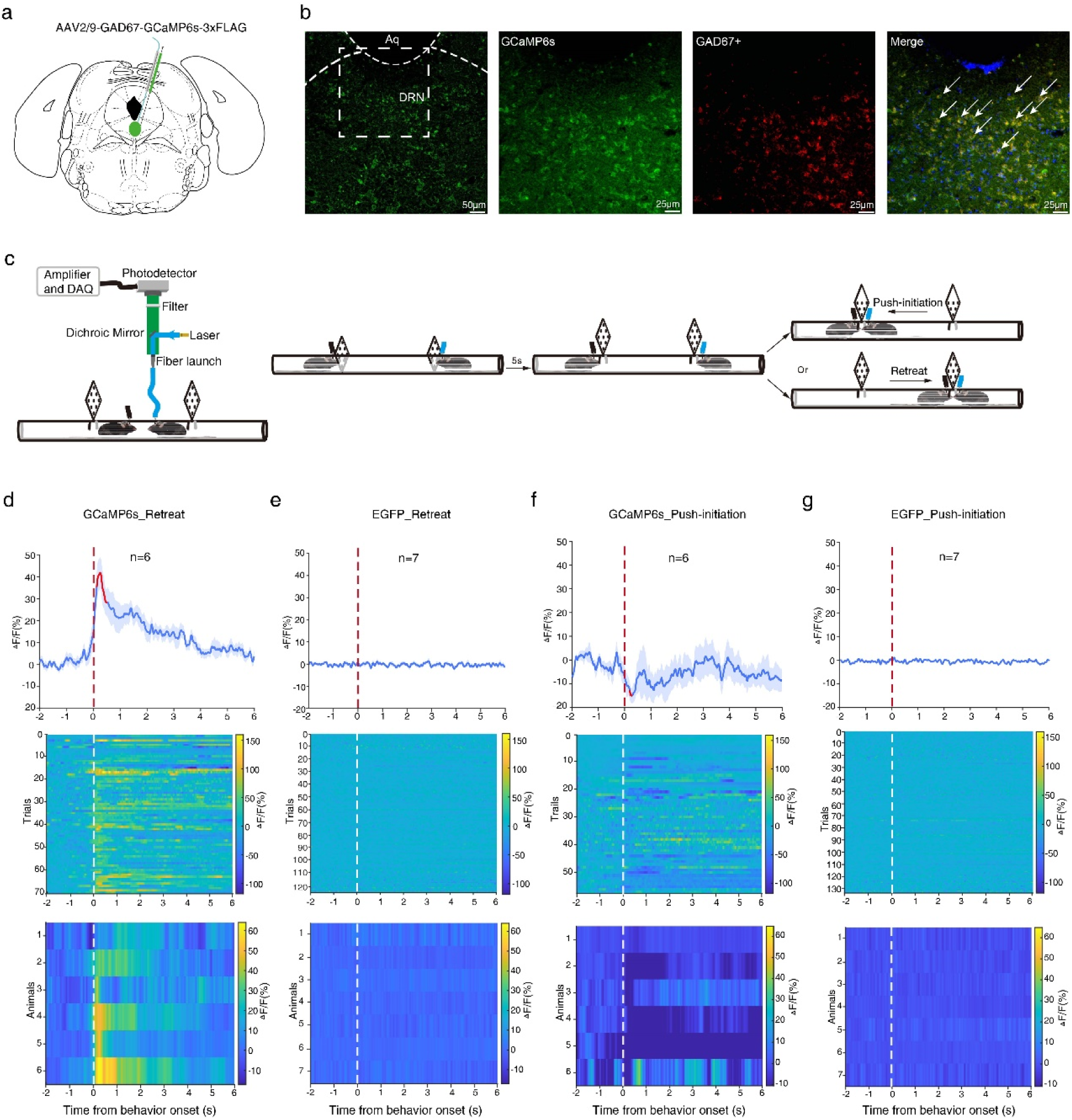
DRN GABAergic neurons were active during retreat but suppressive during push-initiation in the tube test. **a** Schematic representation of the viral injection site in the DRN utilizing AAV2/9-GAD67-GCaMP6s-3xFLAG. **b** Representative photomicrographs of the DRN showing GCaMP6s (green, left) with the enlarged pictures for the boxed area illustrating GCaMP6s expression (green, second subplot), GAD67^+^ (red, third subplot), and their co-localization with DAPI (right). **c** Schematic of the fiber photometry setup (left) and a simplified diagram of the tube test procedure (right). **d** Ca^2+^ signals aligned to the onset of retreats when DRN-GCaMp6f mice meet with their cage mates during the tube test. Upper panel, mean (blue trace) ± SEM (gray shading) showing the average Ca^2+^ signal transients for all trials (n = 6 mice). The red segment indicates a statistically significant increase from baseline (*p* < 0.05; permutation test). Middle panel, the heatmap of Ca^2+^ signals for all trials with each row representing an individual trial (n = 70 trials from 6 mice). Lower panel, the heatmap of Ca^2+^ signals averaged across all trials for each animal (n = 6 mice). **e** Ca^2+^ signals aligned to the onset of retreats in DRN-EGFP mice meeting with their cage mates in the tube test (n = 124 trials from 7 mice). The three panels exhibit similarities to those in panels **d**. **f** Ca^2+^ signals aligned to the onset of push-initiations in DRN-GCaMp6f mice meeting with their cage mates in the tube test (n = 57 trials from 6 mice). The red segment indicates a statistically significant decrease from baseline (*p* < 0.05; permutation test). The three panels show resemblance to those depicted in panels **d**. **g** Ca^2+^ signals aligned to the onset of push-initiations in DRN-EGFP mice meeting with their cage mates in the tube test (n = 133 trials from 6 mice). The three panels exhibit similarities to those in panels **d**.

### Optogenetic activation of DRN GABAergic neurons resulted in loss in the tube test

To elucidate the role of DRN GABAergic neurons in regulating social hierarchy, we employed optogenetic activation of these neurons to monitor real-time changes in the social rank of mice during the tube test. Specifically, the viral vector AAV2/9-GAD67-hChR2(H134R)-EGFP-3xFLAG-WPRE was injected into the DRN (designated as DRN-ChR2 mice; Fig. 2a-b). Subsequently, a fiber optic cannula was implanted 500 μm above the injection site and affixed to the skull. At least four weeks post-injection, we stimulated the DRN in vivo to activate GABAergic neurons using pulses of 473 nm blue light (20 ms, 20 Hz, and 9-19 mW) before the mice entered the tube, maintaining light exposure throughout the duration of the test. Following this stimulation, the mice exhibited a loss of dominance when faced with previously subdominant opponents. Notably, the significant reduction in tube-test ranks persisted for a minimum of two days (Figs. 2c, d, f). Additionally, the optogenetic stimulation led to increased behaviors characterized by push-back, stillness, and retreat, in comparison to pre-photostimulation behaviors (Fig. 2g). However, optogenetic activation of DRN GABAergic neurons did not significantly influence locomotion, anxiety, aggression, social memory, or grip strength (Supplementary Fig. 2). Furthermore, the same photostimulation protocol did not influence the tube test ranks of mice injected with the AAV2/9-GAD67-mCherry-WPRE-hGH-pA viral vector into the DRN (designated as DRN-mCherry mice; Fig. 2e-f), as well as the behaviors in the tube test (Supplementary Fig. 3). Collectively, these findings suggest that optogenetic activation of DRN GABAergic neurons in mice results in a reduction of social dominance, a phenomenon that can be sustained to a certain extent.

**Figure 2.**
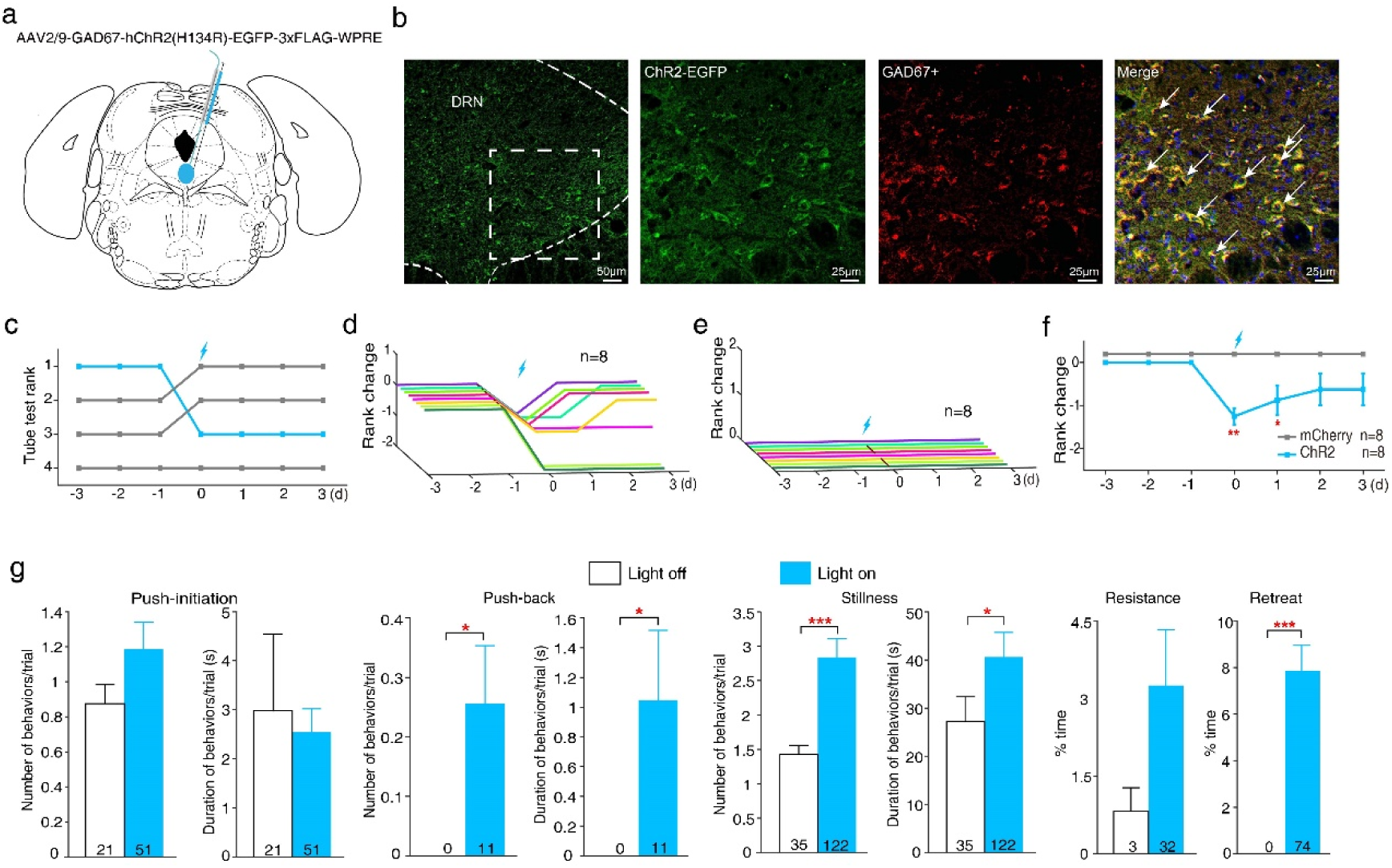
Optogenetic activation of DRN GABAergic neurons led to loss in the tube test. **a** Schematic depiction of the injection site in the DRN for the virus AAV2/9-GAD67-hChR2(H134R)-EGFP-3xFLAG-WPRE. **b** Representative photomicrographs of the DRN depicting ChR2 (green, left) with the enlarged pictures for the boxed area showing ChR2 expression (green, second subplot), GAD67^+^ (red, third subplot), and their co-localization with DAPI (right). **c** Example of the rank positions of a single cohort of mice tested daily over 7 days, indicating that the first-ranked mouse injected with the ChR2 virus fell to the third place following photostimulation. **d** Summary of rank changes in DRN-ChR2 mice before and after the optogenetic activation of DRN GABAergic neurons. Each line represents an individual animal. **e** No rank changes were observed in control mice before and after photostimulation. **f** Comparison of average rank changes in DRN-ChR2 mice following the optogenetic activation of DRN GABAergic neurons delivered throughout the tube test on Day 0 (Wilcoxon signed rank test; Day 0: Z_6_=-2.640, *p*=0.008; Day 1: Z_6_=-2.070, *p*=0.038; Day 2: Z_6_=-1.633, *p*=0.102; Day 3: Z_6_=-1.633, *p*=0.102). **g** Optogenetic activation of DRN GABAergic neurons significantly increased the number (Mann-Whitney U-test: U=432, *p*=0.038) and duration (U=432, *p*=0.039) of push-back, as well as the number (U=223.5, *p*<0.001) and duration (U=362, *p*=0.044) of stillness, and the percentage of time spent retreating (U=60, *p*<0.001). However, no significant differences were observed in the number (U=445, *p*=0.307) and duration (U=415.5, *p*=0.186) of push-initiation or the percentage of time spent resisting (U=424, *p*=0.108). \**p* < 0.05, \*\**p* < 0.01, \*\*\**p*<0.001. Data are expressed as mean ± SEM.

### Chemogenetic activation of DRN GABAergic neurons led to loss in both the tube test and the warm spot test

To further validate the hypothesis that the activation of DRN GABAergic neurons leads to a sustained reduction in social hierarchy among mice, we employed DREADD (designer receptors exclusively activated by designer drugs) technology, which allows for longer modulation periods and a greater number of manipulated neurons. We then extended the findings from the tube test to the warm spot test. The viral vector AAV2/9-GAD67-hM3D(Gq)-mCherry-WPRE was injected into the DRN (referred as DRN- hM3Dq mice; Fig. 3a-b). Four weeks post-injection, Clozapine-N-oxide (CNO, 10 mg/kg) was administered intraperitoneally to one mouse in each cage, while its cage mates received an equivalent volume of saline injection. The tube test was conducted at 1-1.5, 3-5, 6-8, 24, 48, and 72 hours post-injection. Compared to controls, CNO administration resulted in a significant decrease in tube test rankings among the CNO-injected mice, with the most pronounced effect observed between 3 and 5 hours after injection (Fig. 3c-e). CNO-injected mice exhibited a marked increase in passive behaviors, including immobility and retreat (Fig. 3f-g). At 24 hours post-CNO injection, most mice returned to their original ranking positions (Fig. 3d-e). In the warm spot test, DRN-hM3Dq mice exhibited a significant decrease in the duration of warm spot occupancy at two hours post-CNO injection when compared to saline-injected control subjects (Fig. 3h-j). Furthermore, DRN-mCherry mice showed no significant changes in social rank following either CNO or saline injection (Supplementary Fig. 4a-d). Thus, the chemogenetic activation of DRN GABAergic neurons resulted in a reduction of social rank among mice, a finding that can be generalized across different contexts. Additionally, chemogenetic activation was associated with reduced locomotion in the mice (Supplementary Fig. 4e-f).

**Figure 3.**
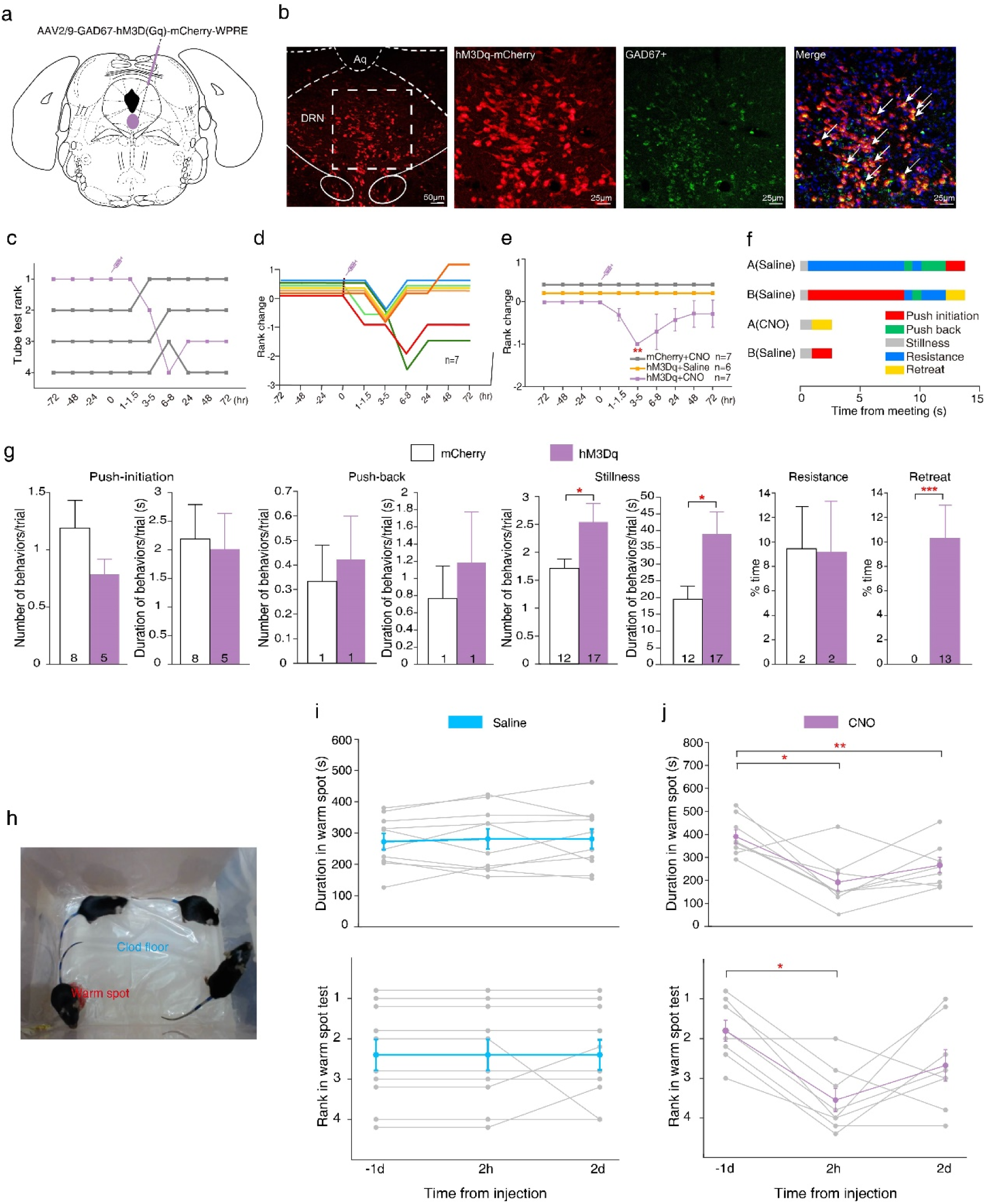
Chemogenetic activation of DRN GABAergic neurons resulted in a loss in both the tube test and the warm spot test. **a** Schematic representation of the injection site in the DRN for the virus AAV2/9-GAD67-hM3D(Gq)-mCherry-WPRE. **b** Representative photomicrographs of the DRN depicting hM3Dq (red, left) with the enlarged pictures for the boxed area showing hM3Dq expression (red, second subplot), GAD67^+^ (green, third subplot), and their co-localization with DAPI (right). **c** Example of the ranking positions for a cage of mice tested daily over a period of 7 days, indicating that the first-ranked mouse injected with the hM3Dq virus was relegated to the last position 6 hours post-CNO injection. **d** Summary of rank changes in DRN-hM3Dq mice before and after CNO injection. Each line represents an individual animal. **e** Difference in average rank change for DRN-hM3Dq mice following CNO injection (Wilcoxon signed rank test; at 1-1.5 hr: Z_5_=-1.414, *p*=0.157; at 3-5 hr: Z_5_=-2.646, *p*=0.008; at 6-8 hr: Z_5_=-1.342, *p*=0.180; at 24 hr: Z_5_=-1.342, *p*=0.180; at 48 hr: Z_5_=-0.816, *p*=0.414; at 72 hr: Z_5_=-0.816, *p*=0.414). **f** Behavioral annotations of two tube test trials between the same pair of mice before and after CNO injection. **g** Chemogenetic activation of DRN GABAergic neurons significantly increased both the number (Mann-Whitney U-test: U=90, *p*=0.043) and duration (U=86, *p*=0.040) of stillness, as well as the percentage of time spent retreating (U=2, *p*<0.001). However, no significant changes were observed in the number (U=129, *p*=0.512) and duration (U=142, *p*=0.864) of push-initiation, the number (U=134, *p*=0.551) and duration (U=133, *p*=0.522) of push-back, or the percentage of time for resistance (U=146, *p*=0.968). **h** Picture for the warm spot test, wherein four mice competed for occupancy of a warm corner situated on an ice-cold floor. **i** Duration of warm spot occupancy (top; One-way repeated measures ANOVA with Bonferroni correction: F_2,18_=0.142; *p*=0.868; n=10) and rank in the warm spot test (bottom; Friedman test with Wilcoxon signed- rank test for multiple comparisons: χ^2^=0.677; *p*=0.717; n=10) for DRN-hM3Dq mice are reported for 1 day before, 2 hours after, and 2 days after saline injection. Each gray folded line represents an individual animal, while the colored line indicates the mean. Animals of the same rank are arranged in parallel to facilitate the presentation of results. **j** Duration of warm spot occupancy (top; F_2,14_=11.785; *p*=0.007; n=8) and rank in the warm spot test (bottom; χ^2^=8.538; *p*=0.014; n=8) for DRN-hM3Dq mice are presented for 1 day before, 2 hours after, and 2 days after CNO injection. \**p*<0.05, \*\**p*<0.01, \*\*\**p*<0.001. Data are expressed as mean ± SEM.

### Optogenetic inhibition of DRN GABAergic neurons induced win in the tube test

To ascertain the necessity of DRN GABAergic neurons in modulating social hierarchy, we employed an optogenetic strategy to selectively inhibit these neurons and subsequently assess changes in social rank among mice during the tube test. Specifically, we administered the viral vector AAV2/9-GAD67-eNpHR3.0-EGFP-WPREs into the DRN (hereafter referred to as DRN-NpHR mice; see Fig. 4a-b). Following this, a fiber optic cannula was implanted 500 μm above the viral injection site and affixed to the skull. At least four weeks post-injection, we stimulated the DRN in vivo to inhibit GABAergic neurons using 589 nm continuous orange light (9-19 mW) immediately prior to the mice entering the tube. The light remained activation throughout the duration of the test. In DRN-NpHR mice, this photostimulation protocol resulted in immediate victories over previously dominant opponents (Fig. 4c-d, f). Conversely, the same protocol did not alter the tube test rankings in DRN-mCherry mice (Fig. 4e-f). Additionally, optogenetic inhibition of DRN GABAergic neurons in subordinate DRN- NpHR mice led to a marked increase in inactivity and a reduction in retreat behaviors compared to the pre-photostimulation condition (Fig. 4g). Importantly, this inhibition did not induce significant changes in anxiety, aggression, social preference, or grip strength in DRN-NpHR mice (Supplementary Fig. 5c-f), as well as the behaviors of DRN-mCherry mice in the tube test (Supplementary Fig. 6). However, it did result in decreased locomotion among the mice (Supplementary Fig. 5b).

**Figure 4.**
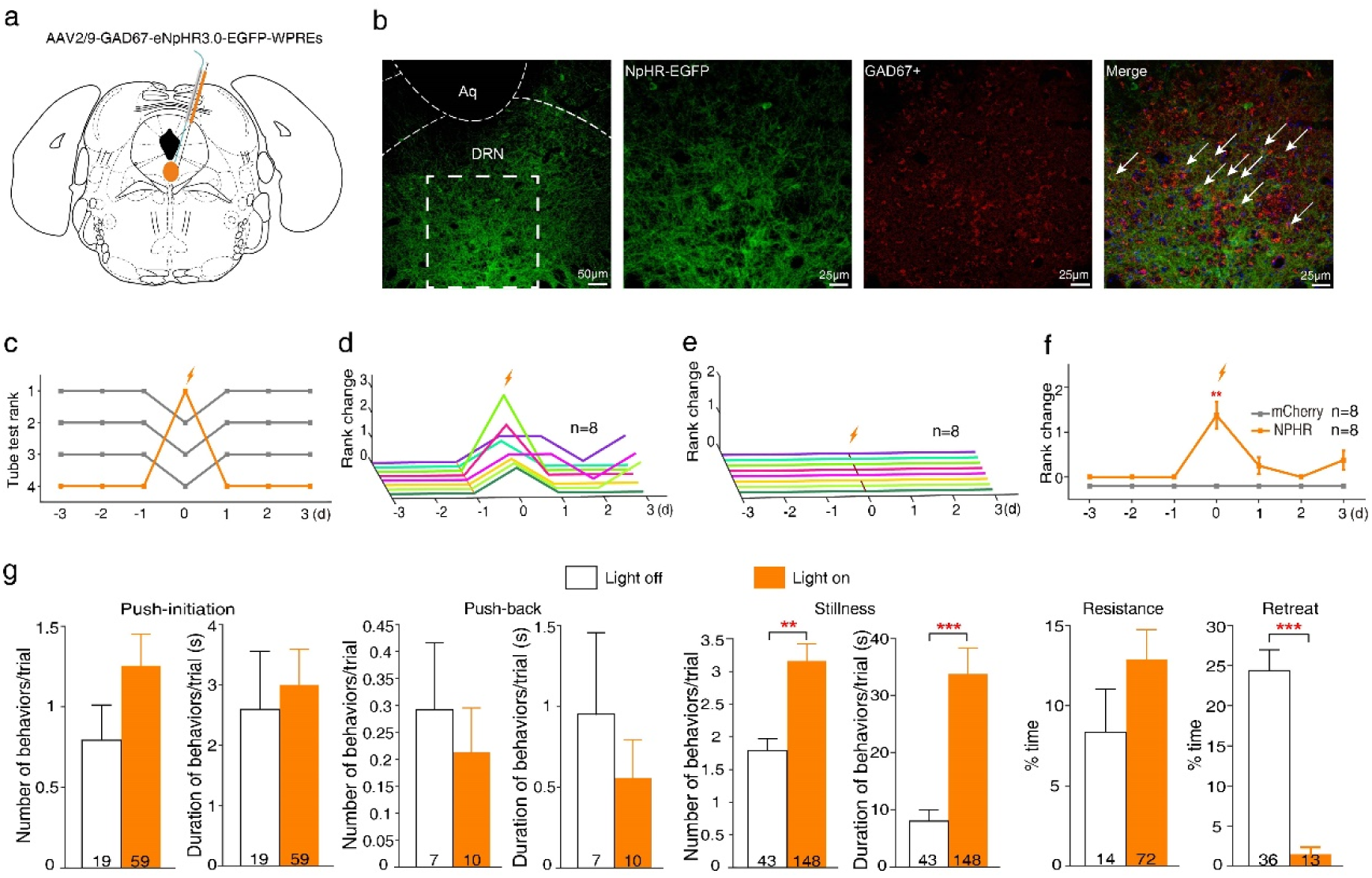
Optogenetic inhibition of DRN GABAergic neurons induced winning in the tube test. **a** Schematic representation of the injection site in the DRN for the virus AAV2/9-GAD67-eNpHR3.0-EGFP-WPREs. **b** Representative photomicrographs of the DRN depicting eNpHR (green, left) with the enlarged pictures for the boxed area showing eNpHR expression (green, second subplot), GAD67^+^ (red, third subplot), and their co-localization with DAPI (right). **c** Example of the ranking positions of a cohort of mice tested daily over 7 days, demonstrating that the mouse initially ranked lowest, which received the eNpHR virus injection, subsequently ascended to the highest rank following photostimulation. **d** Summary of rank changes in DRN-eNpHR mice before and after photostimulation. Each line represents an individual animal. **e** No rank changes were observed in DRN-mCherry mice before and after photostimulation. **f** The average rank change for DRN-eNpHR mice following optogenetic inhibition of DRN GABAergic neurons, administered on Day 0 (Wilcoxon signed rank test; Day 0: Z_6_=- 2.636, *p*=0.008; Day 1: Z_6_=-1.414, *p*=0.157; Day 2: Z_6_=0, *p*=1; Day 3: Z_6_=-1.732, *p*=0.083). **g** Optogenetic inhibition of DRN GABAergic neurons significantly increased the number (Mann-Whitney U test: U=301.5, *p*=0.001) and duration (U=166, *p*<0.001) of stillness, along with the percentage of time dedicated retreat (U=17, *p*<0.001). In contrast, there were no significant changes observed in the number (U=457, *p*=0.168) and duration (U=506.5, *p*=0.463) of push-initiation, the number (U=529.5, *p*=0.520) and duration (U=532, *p*=0.551) of push-back, or the percentage of time spent in resistance (U=428.5, *p*=0.089). \*\**p* < 0.01, \*\*\**p*<0.001. Data are expressed as mean ± SEM.

### Chemogenetic inhibition of DRN GABAergic neurons resulted in win in both the tube test and the warm spot test

To further validate the preceding findings, we employed chemogenetic inhibition to inactivate DRN GABAergic neurons and examined the alterations in social rank among mice during both the tube test and the warm spot test. Specifically, the virus AAV2/9-GAD67-hM4D(Gi)-mCherry-WPREs was administered into the DRN (referred as DRN-hM4Di mice; Fig. 5a-b). Four weeks post-injection of the viral vector, CNO (10 mg/kg) was administered intraperitoneally to one mouse in each cage, while its cage mates received saline injections. The tube test was conducted at 1-1.5, 3-5, 6- 8, 24, 48, and 72 hours following CNO administration (Fig. 5c). Compared to controls, CNO treatment resulted in a significant increase in the tube test rankings of the CNO-injected mice, with the earliest rise observed between 1 and 1.5 hours post-injection, peaking between 3 and 8 hours post-injection (Fig. 5d-e). Notably, there was a significant increase in the frequency of push-initiation, push-back, and resistance, alongside a decrease in the frequency of retreat following CNO administration (Fig. 5f- g). By 24 hours post-injection, most mice returned to their baseline rank positions (Fig. 5c-e). In the warm spot test, DRN-hM4Di mice injected with CNO for two hours demonstrated a significant increase in the duration spent in the warm spot, as well as an elevation in warm spot rank (Fig. 5i), while saline injection did not elicit a comparable change (Fig. 5h). In addition, chemogenetic inhibition did not affect locomotion in mice (Supplementary Fig. 4g-h). Consequently, the inhibition of DRN GABAergic neurons was found to enhance social hierarchy, a change in rank that remained consistent throughout the warm spot test.

**Figure 5.**
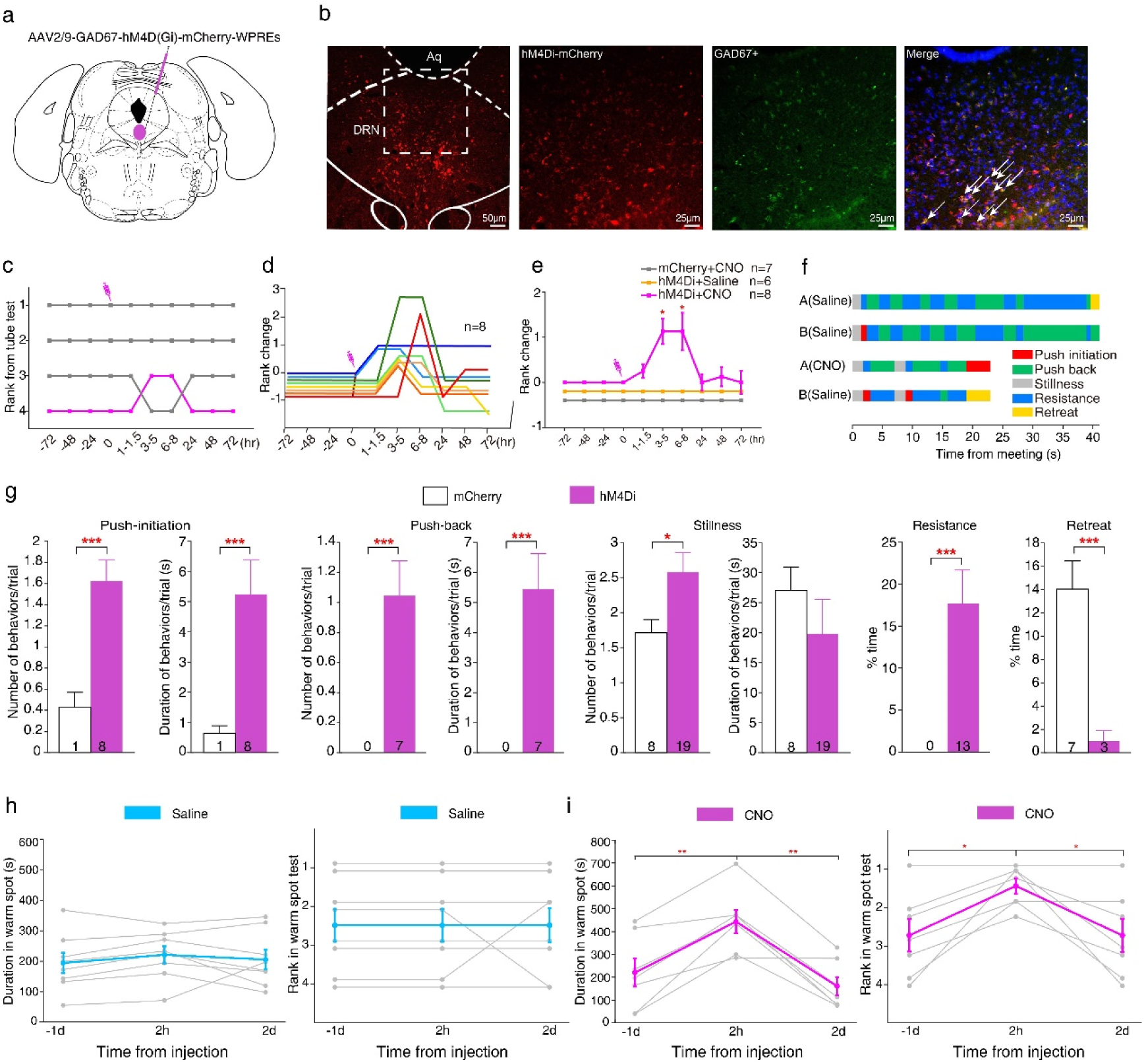
Chemogenetic inhibition of DRN GABAergic neurons produced a win in both the tube test and the warm spot test. **a** Schematic representation of the injection site in the DRN for the virus AAV2/9-GAD67-hM4D(Gi)-mCherry-WPREs. **b** Representative photomicrographs of the DRN depicting hM4Di (red, left) with the enlarged pictures for the boxed area showing hM4Di expression (red, second subplot), GAD67^+^ (green, third subplot), and their co-localization with DAPI (right). **c** Example of the ranking positions of a cohort of mice tested daily over 7 days, demonstrating that the last-ranked mouse injected with the hM4Di virus ascended to third place following CNO administration. **d** Summary of rank changes in DRN-hM4Di mice before and after CNO injection. Each line represents an individual animal. **e** Statistical analysis of mean rank change in DRN-hM4Di mice following CNO injection at 0 hours (Wilcoxon Signed Rank Test; at 3-5 hr: Z_6_=-2.530, *p*=0.011; at 6-8 hr: Z_6_=-2.070, *p*=0.038). **f** Behavioral annotations of two tube test trials between the same pair of mice before and after CNO injection. **g** Chemogenetic inhibition of DRN GABAergic neurons significantly increased the number (Mann-Whitney U-test: U=64.5, *p*<0.001) and duration (U=41.5, *p*<0.001) of push-initiation, the number (U=84, *p*<0.001) and duration (U=84, *p*<0.001) of push-back, the percentage of time spent in resistance (U=51, *p*<0.001), and the number (U=117, *p*=0.011) but not the duration (U=135, *p*=0.050) of stillness. Moreover, the percentage of time spent in retreat was significantly decreased by chemogenetic inhibition (U=18, *p*<0.001). **h** Duration of occupying the warm spot (left; one-way repeated measures ANOVA with Bonferroni correction: F_2,14_=0.769; *p*=0.482; n=8) and rank in the warm spot test (right; Friedman test with Wilcoxon signed-rank test for multiple comparison: χ^2^=0; *p*=1; n=8) for DRN-hM4Di mice assessed at 1 day before, 2 hours after, and 2 days after saline injection. Each gray folded line represents an individual animal, while the colored line represents the average. Mice with the same rank are arranged in parallel to facilitate result presentation. **i** Duration of occupying the warm spot (left; F_2,12_=20.018; *p*<0.001; n=7) and rank in the warm spot test (right; χ^2^=9.579; *p*=0.008; n=7) for DRN-hM4Di mice at 1 day before, 2 hours after, and 2 days after CNO injection. \**p* < 0.05, \*\**p* < 0.01, \*\*\**p*<0.001. Data are expressed as mean ± SEM.

## Discussion

The objective of this study was to elucidate the potential role of DRN GABAergic neurons in the regulation of social hierarchy. To achieve this, we employed optogenetic and chemogenetic techniques in conjunction with fiber photometry recordings. Our findings demonstrate that DRN GABAergic neurons exhibited activation pattern during withdrawal and showed reduced activity during the initiation of push behaviors. Furthermore, both optogenetic and chemogenetic activation of DRN GABAergic neurons resulted in a reduction of social rank, whereas inhibition of these neurons led to a corresponding increase in social rank. These results underscore the bidirectional regulatory role of DRN GABAergic neurons in the context of social hierarchy.

The present findings indicate that DRN plays a pivotal role in regulating social hierarchy, aligning with its established functions such as social interaction, reward processing, arousal, and the modulation of stress and anxiety^22, 26, 36, 42^. The DRN, situated in the dorsal midbrain, encompasses at least four principal cell types that release various neurotransmitters, including serotonin, GABA, glutamate, and dopamine^47, 48^, and serves as the primary source of serotonin within the brain^12, 16, 17^. Previous studies have established a close association between the serotonergic system and social rewards, as well as the acquisition of social dominance across diverse species^9, 29–38^. For instance, the administration of 5-HT into the hemolymph of both lobsters and crayfish reduces the likelihood of retreat and extends the duration of aggressive encounters^18, 39^. Similarly, the administration of 5-HT facilitates the attainment of dominance in both monkeys^34^ and humans^41^. Furthermore, dopamine within various brain regions, including the DRN, is implicated in the representation of social rank signals in green anole lizards (*Anolis carolinensis*)^49^. Collectively, these findings underscore the critical involvement of the DRN and its associated neurotransmitters in the regulation of social hierarchy.

Our findings suggest that GABAergic neurons in the DRN play a critical role in the regulation of social hierarchy. Specifically, activation of DRN GABAergic neurons was associated with an increased frequency of retreat behaviors, whereas inhibition of these neurons resulted in a decreased frequency of retreat in mice. These observations are consistent with previous studies indicating that optogenetic silencing of DRN GABAergic neurons leads to disinhibition of DRN 5-HT neurons, ultimately rescuing social avoidance behavior in mice^24^. GABA, the predominant inhibitory neurotransmitter in the central nervous system, is essential for determining the timing of neuronal firing and maintaining the balance between excitation and inhibition for information processing^50–54^. In the mammalian cerebral cortex, GABAergic interneurons are crucial for regulating this balance and dynamic organization of pyramidal neuron ensembles, which mediate diverse streams of information processing and output channels^55–57^. The connective projections between excitatory and inhibitory neurons within brain regions form neural microcircuits that regulate various social behaviors, including social dominance, social fear, exploration, memory, and preference^58^.

Consistent with this, serotonergic activity relies on a delicate balance between excitatory and inhibitory inputs to DRN 5-HT neurons^42^. Dynamic modulation of 5-HT neurons may be achieved through a push-pull mechanism (convergent excitation and inhibition) or feedforward inhibition provided by local GABAergic neurons^17, 27, 59, 60^. For example, it has been demonstrated that local GABAergic inhibition of 5-HT neurons is fivefold stronger than that induced by external GABAergic afferents^27^, suggesting that DRN GABAergic interneurons exert a robust inhibitory influence on serotonergic firing^24^. This inhibition could diminish 5-HT production by DRN 5-HT neurons, leading to significant reductions in 5-HT levels in the PFC, hippocampus, and hypothalamus, ultimately influencing animal behaviors^27, 42, 61, 62^. Research confirms that DRN 5-HT neurons in mice are activated by reward-related stimuli such as sucrose, food, and social interaction, while DRN GABAergic neurons respond to aversive stimuli including quinine and electrical stimulation, suggesting a functionally antagonistic relationship between these two neuronal types^36^. Together, these findings underscore the potential for GABAergic neurons to serve a complementary role in behavioral modulation relative to 5-HT neurons. Our current results indicate that both optogenetic and chemogenetic activation of DRN GABAergic neurons lead to a reduction in social rank, while inhibition of these neurons corresponds with an increase in social rank. These findings, which contrast with the effects observed when manipulating 5-HT neurons, further highlight the antagonistic interactions between GABAergic and 5-HT neurons in the regulation of social hierarchy.

In summary, our study highlights the critical role of GABAergic neurons in the DRN in regulating social dominance and status. Given the close reciprocal projection relationship between GABAergic and 5-HT neurons in the DRN^42^, we hypothesize that these neuron types may form a microcircuit that modulates behavioral outcomes. Furthermore, GABAergic neurons receive inputs from various upstream targets and connect with multiple cell types within the DRN, influencing relevant behaviors through both direct and indirect pathways. Future research is warranted to elucidate the neural circuits linking DRN GABAergic neurons to their upstream targets, as well as the local neural microcircuits involving interactions between GABAergic and 5-HT or dopaminergic neurons in the DRN that regulate social hierarchy. Moreover, given that GABAergic neurons can be classified into multiple subtypes, it remains to be determined which specific subtype is involved in the regulation of social dominance and associated behaviors.

## Methods

### Animals

Male C57BL/6 mice, aged 8 to 12 weeks, were obtained from Chengdu Dossy Experimental Animals Co., Ltd. The mice were housed in an animal facility maintained at a controlled room temperature of 22 ± 1 °C, with a relative humidity of 60 ± 5%, and were subjected to a 12-hour light/dark cycle (lights on from 08:00 to 20:00). They had ad libitum access to food and water. All experimental procedures were approved by the Animal Care and Use Committee of Chengdu Institute of Biology, Chinese Academy of Sciences (permit number: CIBDWLL2020001), ensuring compliance with established ethical standards.

### Virus injection and fiber implantation

The following viruses were utilized in this study: AAV2/9-GAD67-GCaMP6s- 3xFLAG, AAV2/9-GAD67-hChR2(H134R)-EGFP-3xFLAG-WPRE, AAV2/9- GAD67-eNpHR3.0-EGFP-WPREs, AAV2/9-GAD67-hM3D(Gq)-mCherry-WPRE, AAV2/9-GAD67-hM4D(Gi)-mCherry-WPREs, AAV2/9-GAD67-EGFP-tWPA, and AAV2/9-GAD67-mCherry-WPRE-hGH-pA. These viruses were injected into the DRN at an angle of 15° (AP, -1.3 mm; ML, 0 mm; DV, -2.6 mm from lambda). Each injection delivered a volume of 300 nl of the viral solution. For the optogenetic manipulation experiments, the optical fiber was strategically positioned 0.5 mm above the DRN (angle, 15°; AP, -1.3 mm; ML, 0 mm; DV, -2.1 mm from lambda) to target a broader population of neurons. Conversely, for fiber photometry recordings, the optical fiber was implanted directly at the DRN site (angle, 15°; AP, -1.3 mm; ML, 0 mm; DV, -2.6 mm from lambda), facilitating the acquisition of calcium signal recordings from neurons within the targeted brain region.

### Behavioral tests

*Tube test*. The procedures of the tube test are described in detail in a prior study^46^.

Briefly, to establish a stable social hierarchy, four adult male mice were co-housed in a single cage for a minimum duration of two weeks. Following this period, the mice underwent three consecutive days of handling, succeeded by three consecutive days of training in the tube test. An acrylic tube, measuring 3 cm in inner diameter and 30 cm in length, was utilized for training, with each animal performing ten entries per day (five from each end). During the tube test, pairs of mice were placed at opposite ends of the tube, meeting in the center. The mouse that exited the tube first was designated as the loser, while its counterpart was regarded as the winner. In instances where neither mouse exited the tube within a two-minute timeframe, the trial was repeated. After each trial, the acrylic tube and the surrounding experimental area were cleaned with bathroom paper moistened in 75% alcohol. Each animal participated in three pairwise interactions daily with its cage mates, resulting in a total of six pairs per cage. The ranks among the four mice were established based on the number of victories attained. Stable ranks were defined as those that remained unchanged over a minimum of three consecutive days. Animal behaviors during the tube test were recorded on video and subjected to frame-by-frame analysis using BORIS video analysis software (V7.12.2, University of Torino)^63^. Following the behavioral classification criteria outlined in the previous study^46^, the observed behaviors within the tube were categorized as push- initiation (actively initiating a push), push-back (retaliating after being pushed), resistance (resisting backward movement when pushed), stillness (remaining stationary aside from sniffing and grooming), and retreat (moving backward after being pushed or voluntarily withdrawing). Each behavioral epoch was annotated using BORIS for subsequent analysis. To mitigate potential bias during the analysis, the scorer was blinded to the ranks and prior experiences of the mice while annotating their behaviors. *Warm spot test*. A sufficient volume of water was introduced into a rectangular plastic container (dimensions: 28 cm × 20 cm × 20 cm), which was subsequently placed in a freezer. Prior to the experimental trials, a plastic film was affixed to the ice surface to maintain a temperature of approximately 0°C throughout the testing period. A heating coil, encased in cardboard, was positioned in one corner of the container to elevate the local temperature to 34°C. This configuration was designed to simulate a warm nest, with a diameter of 5 cm, suitable for accommodating a single adult mouse^64^. Before initiating the experiment, four mice from the same cage were housed in a box with an iced bottom but devoid of a warm nest for a duration of 20 minutes to lower their body temperature. Subsequently, the mice were transferred to the experimental container, which was equipped with ice and a warm nest. Their competitive behaviors for the warm nest were documented over a 20-minute observation period.

The warm spot test was conducted on DRN-hM3Dq and DRN-hM4Di mice. One day after the initial warm spot test, each selected individual received an intraperitoneal injection of CNO at a dosage of 10 mg/kg, while its cage mates were administered an equivalent volume of saline solution. The warm spot test was subsequently repeated at 2 and 48 hours post-CNO injection. We employed the BORIS software to document the timestamps for each mouse’s entry into and exit from the warm nest, allowing us to calculate the total duration of nest occupancy for each individual. In instances where multiple animals occupied the warm nest simultaneously, only the mouse occupying the largest area within the nest was recorded. Furthermore, the individual conducting the behavioral scoring was blinded to the hierarchical status and prior experiences of the mice throughout the annotation process.

*Open field test*. To evaluate the motivation for exploring the central region in a novel environment and to monitor the locomotor activity of the test subjects, each mouse was gently placed in the center of a 40 × 40 cm open field apparatus. Mice activity was recorded under dim lighting conditions during the dark phase for a duration of 10 minutes. The open field test was executed using a behavioral research system (CinePlex, Plexon, USA) for video recording and data acquisition. In optogenetic experiments, blue light (473 nm, 20 Hz, 20 ms, 9 mW; MBL-U-473-100mW, New Industry Optoelectronic Tech., Changchun, China) or continuous orange light (589 nm, 9 mW; MGL-F-589-100mW) was intermittently turned on and off in 1-minute epochs. For chemogenetic experiments, the open field test was conducted 2 hours post-administration of CNO. Tracking software (CinePlex Studio V3.5.0) was employed to quantify the number of entries into the central zone, the duration of time spent in the central zone (center time), and the total distance traveled (locomotion) for each minute. *Elevated plus maze*. The Elevated plus maze (EPM) was utilized to assess anxiety-like behavior in mice, employing a configuration comprising two open arms and two closed arms. Each mouse was placed in the maze’s center, while its locomotor activity was recorded using the aforementioned system under dim light conditions during the dark phase for a duration of 9 minutes. For optogenetic stimulation, blue light (473 nm, 20 Hz, 20 ms, 9 mW) or continuous orange light (589 nm, 9 mW) was intermittently activated and deactivated in 3-minute epochs. The time spent in the open versus closed arms was quantified using CinePlex Studio software.

*Social memory test*. The social memory assessment was conducted using an apparatus measuring 40 × 20 cm, comprising three distinct chambers. Initially, the test mouse was acclimatized in the central chamber for a duration of 10 minutes. Subsequently, an unfamiliar mouse, referred to as the object mouse, was introduced into one of the side chambers. Following a 5-minute familiarization period with the object mouse, a novel object mouse was introduced into the opposing side chamber. During the subsequent 5-minute observation period, the exploration behaviors of the test mice towards the familiar and novel object mice were recorded. For analysis, the time spent by the test mice in the interaction zone (defined as half of the central chamber adjacent to the side chamber, measuring 10 × 20 cm) was quantified in 1-minute epochs. During these epochs, the mice were subjected to alternating light-on and light-off conditions, utilizing 473 nm blue light (20 Hz, 20 ms, 9 mW), or 589 nm continuous orange light (9 mW) that was intermittently activated and deactivated in 1-minute intervals.

*Resident-intruder test*. The standard resident-intruder test was performed on the experimental mice. Prior to the initiation of the test, all cage mates were removed from the home cages, allowing the test subjects to acclimatize to their environment for a duration of 10 minutes. Following this acclimatization period, unfamiliar adolescent male C57BL/6 mice (aged 5-7 weeks) were introduced as intruders for a test period of 10 minutes. During this time, blue light (473 nm, 20 Hz, 20 ms, 9 mW) or continuous orange light (589 nm, 9 mW) was alternately turned on and off in 1-minute epochs. The frequency and duration of aggressive behaviors (such as chasing and attacking) as well as non-aggressive social interactions (including sniffing and social grooming) were meticulously analyzed for each 1-minute epoch, correlating with the light-on and light-off periods.

*Grip-strength measurement*. We employed a force gauge (DS2-50N, Puyan, Dongguan, China) to assess the forelimb grip strength of mice. Each mouse underwent measurements across three experimental blocks, with two-hour intervals between blocks. Within each block, mice participated in 10 trials conducted over approximately 6 minutes, divided into two equal 3-minute periods corresponding to light-on and light-off conditions. The former conditions were illuminated by either blue light (473 nm, 20 Hz, 20 ms, 9 mW) or continuous orange light (589 nm, 9 mW). The maximum grip strengths recorded during both light-on and light-off conditions across the three blocks were subsequently compiled for statistical analysis.

### Fiber photometry recordings

Mice were administered an injection of 300 nl of the AAV2/9-GAD67-GCaMP6s- 3xFLAG virus, followed by the implantation of a fiber optic cannula into the DRN. Following viral expression, fiber photometry recordings were conducted utilizing a recording system (QAXK-FPS-SS-LED, ThinkerTech, Nanjing, China). The GCaMP6s virus was activated with 488 nm excitation light, while signals at 405 nm were subtracted from those at 488 nm to yield corrected signal outputs. The sampling frequency was set to 100 Hz, with the laser intensity maintained at a low level (40 µW) at the tip of the optical fiber to minimize photobleaching. Behavioral observations during the tube test were captured using a camera positioned adjacent to the tube and annotated by BORIS for subsequent analysis. To synchronize the fiber photometry recordings with the video data, an external trigger emitted a TTL pulse to the recording system while simultaneously activating a red light-emitting diode, which was visible to the camera. For each pair of opponents, six trials were conducted, with each side initiating three entries. To establish control measures, mice were also allowed to traverse the tube alone across six trials, with each side initiating three entries.

The data were analyzed utilizing vendor-provided code from ThinkerTech Nanjing Biotech Co., Ltd., implemented in MATLAB. The change in fluorescence, denoted as ΔF/F₀, was calculated using the formula (F - F₀)/F₀, where F₀ represents the baseline fluorescence signal averaged over a one-second interval preceding the initiation of a behavioral epoch. For Ca²⁺ signal analysis, the onset of each behavior was standardized to time zero. In the assessment of control signals during the tube-walking task, fluorescence signals were synchronized to the moment when mice reached the midpoint of the tube. A permutation test was employed to evaluate the statistical significance of fluorescence changes associated with the behaviors, as previously documented^36^.

### Optogenetic manipulation of DRN GABAergic neurons in the tube test

The procedures employed in this study have been delineated in detail in previous studies^46, 64^. Briefly, mice were administered one of the following viruses: AAV2/9- GAD67-hChR2(H134R)-EGFP-3xFLAG-WPRE, AAV2/9-GAD67-eNpHR3.0-EGFP-WPRE, or AAV2/9-GAD67-mCherry-WPRE-hGH-pA, via injection into the DRN, followed by the implantation of a fiber optic cannula. Once a stable tube test rank was established for a duration exceeding three days, all four mice were connected to dummy optic fibers and traversed a tube equipped with a 12 mm slit at the top for two days. Mice with comparatively lower ranks were subjected to optogenetic inhibition, while those ranked higher underwent optogenetic activation. On the test day, the tube test commenced initially without light, followed by the application of photostimulation at 473 nm, 20 Hz, 20 ms (or 589 nm, continuous) prior to the mice entering the test tube. For each subject, photostimulation began at an intensity of 9 mW, with the tube test conducted first against their closest-ranked cage mate, followed by tests against increasingly distanced rank counterparts. If rank changes occurred, the mice exchanged starting positions to repeat the test. Each mouse was required to secure two out of three victories from each side of the tube. A successful inference of rank change due to photostimulation at this specific intensity was determined if the test mice won or lost from both sides of the tube. Otherwise, the light intensity was incrementally increased and the same procedure would be repeated until a rank change was achieved or the light intensity reached 20 mW. Only one mouse from each test cage served as the light-regulated subject, and the total number of victories induced by photostimulation for each test mouse was recorded.

### Chemogenetic manipulation of DRN GABAergic neurons in the tube test

Mice were administered either AAV2/9-GAD67-hM3D(Gq)-mCherry-WPRE, AAV2/9-GAD67-hM4D(Gi)-mCherry-WPRE, or AAV2/9-GAD67-mCherry-WPRE-hGH-pA viral vectors into the DRN. Following a minimum four-week recovery period post-injection, all four mice housed together received an intraperitoneal (i.p.) injection of saline. A tube test was subsequently conducted at 1-1.5, 3-5, 6-8, 24, 48, and 72 hours post-injection. One week later, the test mice were administered an i.p. injection of CNO (10 mg/kg), while the control group received saline, and the tube test was repeated at the same aforementioned time intervals. Mice were randomly selected as test subjects. Statistical analyses were performed to evaluate rank changes for each mouse.

### Immunofluorescence

Upon completion of all experimental procedures, mice were perfused with phosphate-buffered saline (PBS), followed by 4% paraformaldehyde (PFA). Following dissection, the brains were post-fixed overnight in 4% PFA and subsequently subjected to a dehydration protocol involving 15% sucrose in PFA for 24 hours, followed by an additional 24 hours in 30% sucrose in PFA. Coronal brain sections, each 30 μm thick, were serially prepared using a cryostat (CM1850, LEICA, Germany). The primary antibody, rabbit monoclonal anti-GAD67 (1:1000, ab213508, Abcam), was incubated overnight at 4°C, after which sections were washed with PBS. Subsequently, Alexa-488 or Alexa-647 conjugated secondary antibodies, namely Donkey Anti-Rabbit IgG H&L (Alexa Fluor® 488) (ab150061, Abcam) and Donkey Anti-Rabbit IgG H&L (Alexa Fluor® 647) (ab150063, Abcam), were applied at a dilution of 1:500 for 2 hours at room temperature. Following this treatment, cell nuclei were stained with DAPI for 5 minutes. Images of the stained sections were captured using a fluorescence microscope (Nexcope NE910, Ningbo, China).

### Statistical analysis and reproducibility

The normality of distributions and the homogeneity of variances for all values were assessed using the Shapiro-Wilk test and Levene’s test, respectively. In the tube test, the Mann-Whitney U test was employed to evaluate differences in behavioral parameters across various light conditions and among distinct groups. For the warm spot test, one-way repeated measures ANOVA with Bonferroni correction was utilized to compare the duration of occupancy in the warm spot, while the Friedman test was applied to assess rank differences across multiple time points. In the open field test, one-way repeated measures ANOVA with Bonferroni correction was used to analyze behavioral parameter differences among groups subjected to chemogenetic manipulation. Additionally, paired-samples t-tests were conducted to compare behavioral parameters between different light conditions in optogenetic manipulation. For the elevated plus-maze test, a two-way repeated measures ANOVA with Bonferroni correction was utilized, incorporating the factors of "light condition" and "group". In the resident-intruder test, the Wilcoxon signed-rank test was utilized. For the social memory test, a two-way repeated measures ANOVA with Bonferroni correction was conducted, considering the factors of "zone" and "group". Furthermore, paired-samples t-tests were performed to compare forelimb grip strength across different light conditions. Statistical analyses were carried out using SPSS software (version 23.0), with significance levels set at *p* < 0.05 and highly significant levels at *p* < 0.001. The reproducibility of micrographic and behavioral experiments in optogenetics, chemogenetics, and photometry recordings was consistent with the actual number of experimental subjects.

## Reporting summary

Further information on research design is available in the Nature Portfolio Reporting Summary linked to this article.

## Data availability

The dataset generated and analyzed during the current study are available in the GitHub repository: https://github.com/myleafmavis/Data-DRN-GABA

## Code availability

Matlab code for analyzing the data and generating the figures is available at: https://github.com/myleafmavis/code-for-DRN-GABA

## Acknowledgements

This study was supported by the grants from the National Natural Science Foundation of China (Nos. 32170504 and 31970422).

## Author contributions

Y.-Z.F. and G.-Z.F. contributed the design of the experiments. L.-D.L., Y.-Z.F., S.-X.G. and Y.W. performed the surgeries and behavioral tests. L.-D.L. and S.-X.G. analyzed all the data. L.-D.L. and Y.W. performed immunohistochemistry. Z.-Y.W., T.Q. and S.-X.S performed mice husbandry. L.-D.L., Y.-Z.F., and G.-Z.F. wrote the manuscript. L.- D.L., Y.-Z.F., and G.-Z.F. contributed the textual revision of the paper. All authors discussed the paper.

## Competing interests

The authors declare no competing interests.

**Correspondence** and requests for materials should be addressed to Guangzhan Fang.

## References

1. Strauss, E.D., Curley, J.P., Shizuka, D., Hobson, E.A. The centennial of the pecking order: current state and future prospects for the study of dominance hierarchies. Phil. Trans. R. Soc. B 377, 20200432 (2022).

2. Marmot, M. Status syndrome: how your social standing directly affects your health and life expectancy. Br. Med. J. 329, 407–409 (2004).

3. Larrieu, T., Sandi, C. Stress-induced depression: is social rank a predictive risk factor? Bioessays 40, e1800012 (2018).

4. Pusey, A., Williams, J., Goodall, J. The influence of dominance rank on the reproductive success of female chimpanzees. Science 277, 828–831 (1997).

5. Sapolsky, R.M. The influence of social hierarchy on primate health. Science 308, 648–652 (2005).

6. Strauss, E.D., Holekamp, K.E. Social alliances improve rank and fitness in convention-based societies. Proc. Natl. Acad. Sci. U.S.A. 116, 8919–8924 (2019).

7. Milewski, T.M., Lee, W., Champagne, F.A., Curley, J.P. Behavioural and physiological plasticity in social hierarchies. Phil. Trans. R. Soc. B 377, 20200443 (2022).

8. Wang, F., Kessels, H.W., Hu, H.L. The mouse that roared: neural mechanisms of social hierarchy. Trends Neurosci. 37, 674–682 (2014).

9. Watanabe, N., Yamamoto, M. Neural mechanisms of social dominance. Front. Neurosci. 9, 154 (2015).

10. Dwortz, M.F., Curley, J.P., Tye, K.M., Padilla-Coreano, N. Neural systems that facilitate the representation of social rank. Phil. Trans. R. Soc. B 377, e20200444 (2022).

11. Zhou, T.T., Sandi, C., Hu, H.L. Advances in understanding neural mechanisms of social dominance. Curr. Opin. Neurobiol. 49, 99–107 (2018).

12. Warden, M.R., Selimbeyoglu, A., Mirzabekov, J.J., Lo, M., Thompson, K.R., Kim, S.-Y., Adhikari, A., Tye, K.M., Frank, L.M., Deisseroth, K. A prefrontal cortex-brainstem neuronal projection that controls response to behavioural challenge. Nature 492, 428–432 (2012).

13. Mah, L., Arnold, M.C., Grafman, J. Impairment of social perception associated with lesions of the prefrontal cortex. Am. J. Psychiat. 161, 1247–1255 (2004).

14. Fujii, N., Hihara, S., Nagasaka, Y., Iriki, A. Social state representation in prefrontal cortex. Social Neurosci. 4, 73–84 (2009).

15. Wang, F., Zhu, J., Zhu, H., Zhang, Q., Lin, Z.M., Hu, H.L. Bidirectional control of social hierarchy by synaptic efficacy in medial prefrontal cortex. Science 334, 693–697 (2011).

16. Challis, C., Beck, S.G., Berton, O. Optogenetic modulation of descending prefrontocortical inputs to the dorsal raphe bidirectionally bias socioaffective choices after social defeat. Front. Behav. Neurosci. 8, 43 (2014).

17. Geddes, S.D., Assadzada, S., Lemelin, D., Sokolovski, A., Bergeron, R., Haj-Dahmane, S., Beique, J.C. Target-specific modulation of the descending prefrontal cortex inputs to the dorsal raphe nucleus by cannabinoids. Proc. Natl. Acad. Sci. U.S.A. 113, 5429–5434 (2016).

18. Huber, R., Smith, K., Delago, A., Isaksson, K., Kravitz, E.A. Serotonin and aggressive motivation in crustaceans: altering the decision to retreat. Proc. Natl. Acad. Sci. U.S.A. 94, 5939–5942 (1997).

19. Audero, E., Mlinar, B., Baccini, G., Skachokova, Z.K., Corradetti, R., Gross, C. Suppression of serotonin neuron firing increases aggression in mice. J. Neurosci. 33, 8678–8688 (2013).

20. Caramaschi, D., de Boer, S.F., Koolhaas, J.M. Differential role of the 5-HT1A receptor in aggressive and non-aggressive mice: an across-strain comparison. Physiol. Behav. 90, 590–601 (2007).

21. Cools, R., Roberts, A.C., Robbins, T.W. Serotoninergic regulation of emotional and behavioural control processes. Trends Cognit. Sci. 12, 31–40 (2008).

22. Paquelet, G.E., Carrion, K., Lacefield, C.O., Zhou, P., Hen, R., Miller, B.R. Single-cell activity and network properties of dorsal raphe nucleus serotonin neurons during emotionally salient behaviors. Neuron 110, 2664–2679 (2022).

23. Monti, J.M. The role of dorsal raphe nucleus serotonergic and non-serotonergic neurons, and of their receptors, in regulating waking and rapid eye movement (REM) sleep. Sleep Med. Rev. 14, 319–327 (2010).

24. Challis, C., Boulden, J., Veerakumar, A., Espallergues, J., Vassoler, F.M., Pierce, R.C., Beck, S.G., Berton, O. Raphe GABAergic neurons mediate the acquisition of avoidance after social defeat. J. Neurosci. 33, 13978–13988 (2013).

25. Gervasoni, D., Peyron, C., Rampon, C., Barbagli, B., Chouvet, G., Urbain, N., Fort, P., Luppi, P.H. Role and origin of the GABAergic innervation of dorsal raphe serotonergic neurons. J. Neurosci. 20, 4217–4225 (2000).

26. Roche, M., Commons, K.G., Peoples, A., Valentino, R.J. Circuitry underlying regulation of the serotonergic system by swim stress. J. Neurosci. 23, 970–977 (2003).

27. Zhou, L., Liu, M.Z., Li, Q., Deng, J., Mu, D., Sun, Y.G. Organization of functional long- range circuits controlling the activity of serotonergic neurons in the dorsal raphe nucleus. Cell Rep. 18, 3018–3032 (2017).

28. Spoida, K., Masseck, O.A., Deneris, E.S., Herlitze, S. Gq/5-HT2c receptor signals activate a local GABAergic inhibitory feedback circuit to modulate serotonergic firing and anxiety in mice. Proc. Natl. Acad. Sci. U.S.A. 111, 6479–6484 (2014).

29. Larson, E.T., Summers, C.H. Serotonin reverses dominant social status. Behav. Brain Res. 121, 95–102 (2001).

30. Korzan, W.J., Summers, C.H. Serotonergic response to social stress and artificial social sign stimuli during paired interactions between male *Anolis carolinensis*. Neuroscience 123, 835–845 (2004).

31. Cases, O., Seif, I., Grimsby, J., Gaspar, P., Chen, K., Pournin, S., Muller, U., Aguet, M., Babinet, C., Shih, J.C., Demaeyer, E. Aggressive behavior and altered amounts of brain serotonin and norepinephrine in mice lacking MAOA. Science 268, 1763–1766 (1995).

32. Coccaro, E.F. Impulsive aggression and central serotonergic system function in humans: an example of a dimensional brain-behavior relationship. Int. Clin. Psychopharmacol. 7, 3–12 (1992).

33. Yeh, S.R., Fricke, R.A., Edwards, D.H. The effect of social experience on serotonergic modulation of the escape circuit of crayfish. Science 271, 366–369 (1996).

34. Raleigh, M.J., McGuire, M.T., Brammer, G.L., Pollack, D.B., Yuwiler, A. Serotonergic mechanisms promote dominance acquisition in adult male vervet monkeys. Brain Res. 559, 181–190 (1991).

35. Sandi, C., Haller, J. Stress and the social brain: behavioural effects and neurobiological mechanisms. Nat. Rev. Neurosci. 16, 290–304 (2015).

36. Li, Y., Zhong, W.X., Wang, D.Q., Feng, Q.R., Liu, Z.X., Zhou, J.F., Jia, C.Y., Hu, F., Zeng,J.W., Guo, Q.C., Fu, L., Luo, M.M. Serotonin neurons in the dorsal raphe nucleus encode reward signals. Nat. Commun. 7, 1–15 (2016).

37. Terranova, J.I., Song, Z.M., Larkin, T.E., Hardcastle, N., Norvelle, A., Riaz, A., Albers, H.E. Serotonin and arginine-vasopressin mediate sex differences in the regulation of dominance and aggression by the social brain. Proc. Natl. Acad. Sci. U.S.A. 113, 13233–13238 (2016).

38. Edwards, D.H., Kravitz, E. A. Serotonin, social status and aggression. Curr. Opin. Neurobiol. 7, 812–819 (1997).

39. Livingstone, M.S., Harriswarrick, R.M., Kravitz, E.A. Serotonin and octopamine produce opposite postures in lobsters. Science 208, 76–79 (1980).

40. Raleigh, M.J., McGuire, M.T., Brammer, G.L., Yuwiler, A. Social and environmental influences on blood serotonin concentrations in monkeys. Arch. Gen. Psychiatry 41, 405–410 (1984).

41. Moskowitz, D.S., Pinard, G., Zuroff, D.C., Annable, L., Young, S.N. The effect of tryptophan on social interaction in everyday life: A placebo-controlled study. Neuropsychopharmacology 25, 277–289 (2001).

42. Hernandez-Vazquez, F., Garduno, J., Hernandez-Lopez, S. GABAergic modulation of serotonergic neurons in the dorsal raphe nucleus. Rev. Neurosci. 30, 289–303 (2019).

43. Weissbourd, B., Ren, J., DeLoach, K.E., Guenthner, C.J., Miyamichi, K., Luo, L. Presynaptic partners of dorsal raphe serotonergic and GABAergic neurons. Neuron 83, 645–662 (2014).

44. Martın-Ruiz, R., Puig, M.V., Celada, P., Shapiro, D.A., Roth, B.L., Mengod, G., Artigas, F. Control of serotonergic function in medial prefrontal cortex by serotonin-2A receptors through a glutamate-dependent mechanism. J. Neurosci. 21, 9856–9866 (2001).

45. Li, Y., Li, C.Y., Xi, W., Jin, S., Wu, Z.H., Jiang, P., Dong, P., He, X.B., Xu, F.Q., Duan, S., Zhou, Y.D., Li, X.M. Rostral and caudal ventral tegmental area GABAergic Inputs to different dorsal raphe neurons participate in opioid dependence. Neuron 101, 748–761 (2019).

46. Fan, Z.X., Zhu, H., Zhou, T.T., Wang, S., Wu, Y., Hu, H.L. Using the tube test to measure social hierarchy in mice. Nat. Protoc. 14, 819–831 (2019).

47. Hioki, H., Nakamura, H., Ma, Y.F., Konno, M., Hayakawa, T., Nakamura, K.C., Fujiyama, F., Kaneko, T. Vesicular glutamate transporter 3-expressing nonserotonergic projection neurons constitute a subregion in the rat midbrain raphe nuclei. J. Comp. Neurol. 518, 668–686 (2010).

48. Okaty, B.W., Freret, M.E., Rood, B.D., Brust, R.D., Hennessy, M.L., Debairos, D., Kim, J.C., Cook, M.N., Dymecki, S.M. Multi-scale molecular deconstruction of the serotonin neuron system. Neuron 88, 774–791 (2015).

49. Korzan, W.J., Forster, G.L., Watt, M.J., Summers, C.H. Dopaminergic activity modulation via aggression, status, and a visual social signal. Behav. Neurosci. 120, 93–102 (2006).

50. Markram, H., Toledo-Rodriguez, M., Wang, Y., Gupta, A., Silberberg, G., Wu, C.Z. Interneurons of the neocortical inhibitory system. Nat. Rev. Neurosci. 5, 793–807 (2004).

51. Trevelyan, A.J., Sussillo, D., Watson, B.O., Yuste, R. Modular propagation of epileptiform activity: Evidence for an inhibitory veto in neocortex. J. Neurosci. 26, 12447–12455 (2006).

52. Klausberger, T., Somogyi, P. Neuronal diversity and temporal dynamics: the unity of hippocampal circuit operations. Science 321, 53–57 (2008).

53. Haider, B., McCormick, D.A. Rapid neocortical dynamics: cellular and network mechanisms. Neuron 62, 171–189 (2009).

54. Salat, K., Kulig, K., Gajda, J., Wieckowski, K., Filipek, B., Malawska, B. Evaluation of anxiolytic-like, anticonvulsant, antidepressant-like and antinociceptive properties of new 2- substituted 4-hydroxybutanamides with affinity for GABA transporters in mice. Pharmacol. Biochem. Behav. 110, 145–153 (2013).

55. Roux, L., Buzsaki, G. Tasks for inhibitory interneurons in intact brain circuits. Neuropharmacology 88, 10–23 (2015).

56. Harris, K.D., Shepherd, G.M.G. The neocortical circuit: themes and variations. Nat. Neurosci. 18, 170–181 (2015).

57. Huang, Z.J. Toward a genetic dissection of cortical circuits in the mouse. Neuron 83, 1284–1302 (2014).

58. Xu, S., Jiang, M., Liu, X., Sun, Y.H., Yang, L., Yang, Q.H., Bai, Z.T. Neural circuits for social interactions: from microcircuits to input-output circuits. Front. Neural Circuits 15, 768294 (2021).

59. Isaacson, J.S., Scanziani, M. How inhibition shapes cortical activity. Neuron 72, 231–243 (2011).

60. Amo, R., Fredes, F., Kinoshita, M., Aoki, R., Aizawa, H., Agetsuma, M., Aoki, T., Shiraki, T., Kakinuma, H., Matsuda, M., Yamazaki, M., Takahoko, M., Tsuboi, T., Higashijima, S., Miyasaka, N., Koide, T., Yabuki, Y., Yoshihara, Y., Fukai, T., Okamoto, H. The habenulo- raphe serotonergic circuit encodes an aversive expectation value essential for adaptive active avoidance of danger. Neuron 84, 1034–1048 (2014).

61. Xu, Z., Feng, Z., Zhao, M., Sun, Q., Deng, L., Jia, X., Jiang, T., Luo, P., Chen, W., Tudi, A., Yuan, J., Li, X., Gong, H., Luo, Q., Li, A. Whole-brain connectivity atlas of glutamatergic and GABAergic neurons in the mouse dorsal and median raphe nuclei. Elife 10, (2021).

62. Muzerelle, A., Scotto-Lomassese, S., Bernard, J.F., Soiza-Reilly, M., Gaspar, P. Conditional anterograde tracing reveals distinct targeting of individual serotonin cell groups (B5-B9) to the forebrain and brainstem. Brain Struct. Funct. 221, 535–561 (2016).

63. Friard, O., Gamba, M., Fitzjohn, R. BORIS: a free, versatile open-source event-logging software for video/audio coding and live observations. Methods Ecol. Evol. 7, 1325–1330 (2016).

64. Zhou, T.T., Zhu, H., Fan, Z.X., Wang, F., Chen, Y., Liang, H.X., Yang, Z.F., Zhang, L., Lin, L.N., Zhan, Y., Wang, Z., Hu, H.L. History of winning remodels thalamo-PFC circuit to reinforce social dominance. Science 357, 162–168 (2017).

