## Supplementary Figures for "GABAergic neurons in the dorsal raphe nucleus regulate social hierarchy in mice"

**This file includes:**

Figure S1-S6.

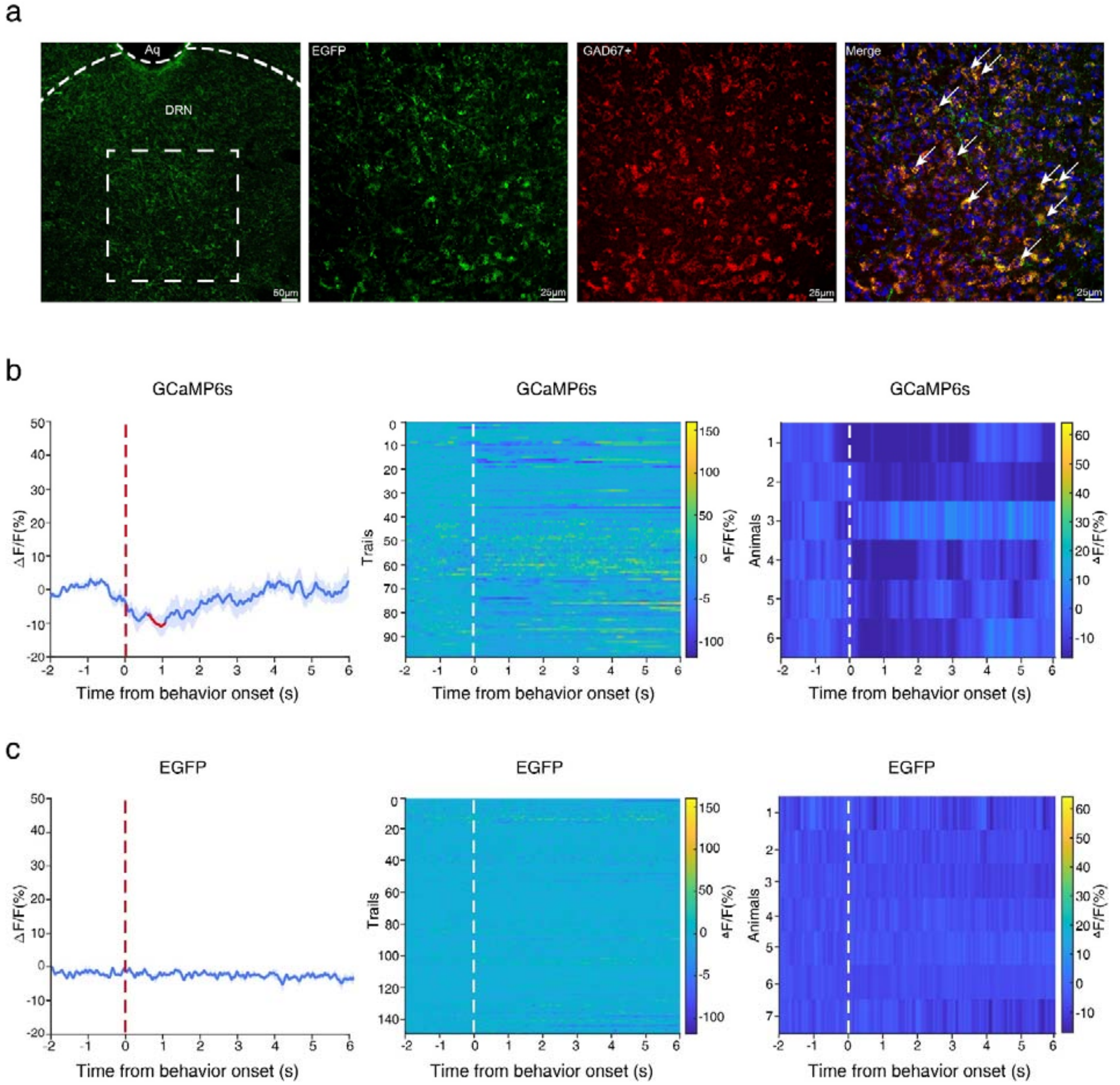


**Figure S1. DRN GABAergic neurons were inhibited during effortful behaviors (including push-initiation, push-back, and resistance) in the tube test. a** Representative photomicrograph of the DRN showing EGFP (green, left) with the enlarged images for the boxed area illustrating EGFP expression (green, second subplot), GAD67$^{+}$ (red, third subplot), and their co-localization with DAPI (right). **b** $Ca^{2+}$ signals aligned to the onset of effortful behaviors when DRN-GCaMp6f mice meet with their cage mates in the tube test. Left panel, mean (blue trace) ± SEM (gray shading) of the average $Ca^{2+}$ signal transients for all trials (n = 6 mice). The red segment indicates a statistically significant increase from baseline ($p < 0.05$; permutation test). Middle panel, the heatmap of $Ca^{2+}$ signals for all trials with each row representing an individual trial (n = 100 trials from 6 mice). Right panel, the heatmap of $Ca^{2+}$ signals for all animals with each row represents the average of $Ca^{2+}$ signals across all trials for each animal (n = 6 mice). **c** $Ca^{2+}$ signals aligned to the onset of effortful behaviors in DRN-EGFP mice meeting with their cage mates during the tube test (n = 149 trials from 7 mice). The three panels exhibit a similarity to those depicted in panels **b**.

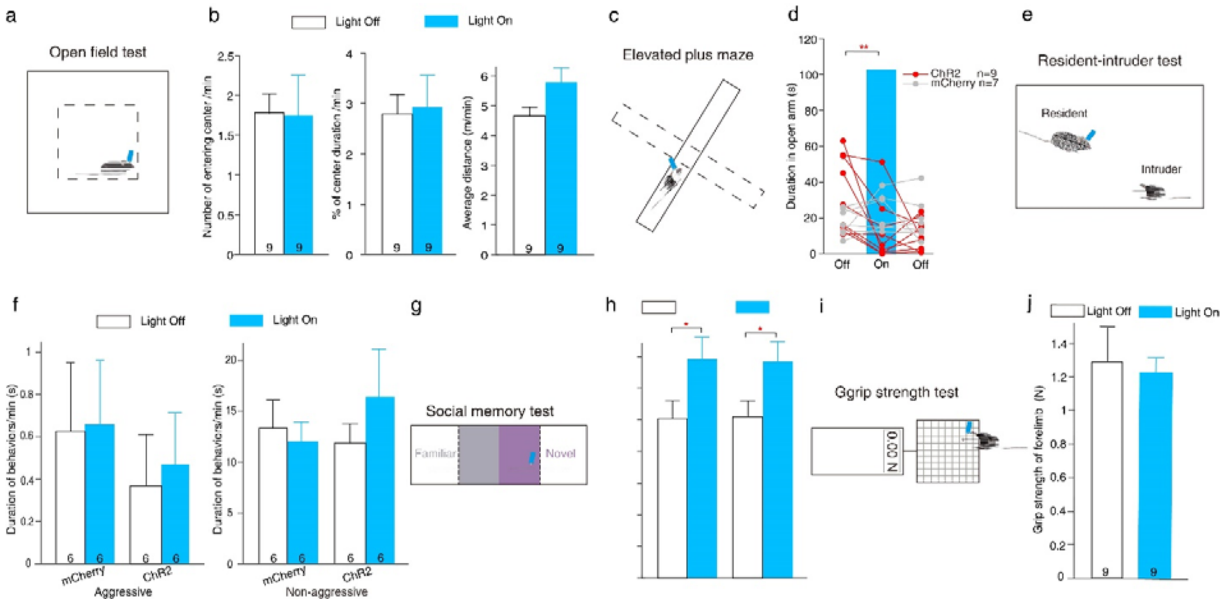


**Figure S2. Optogenetic activation of DRN GABAergic neurons did not significantly affect locomotion, aggression, social memory, or grip strength in mice. a** Schematics representation of the open field test (OFT). Blue light (473 nm, 20 Hz, 20 ms, and 9 mW) was intermittently activated and deactivated for 1-minute epochs over a duration of 10 minutes. **b** Average number of entries into the center (left; Paired samples t-test; $t_8$=0.367, *p*=0.723), percentage of time spent in the center (middle; $t_8$=-0.338, *p*=0.744), and average total distance traveled (right; $t_8$=-2.079, *p*=0.071). **c** Schematics representation of the elevated plus maze. Blue light was intermittently turned on and turned off for 3-minute epochs over a duration of 9 minutes. **d** Duration spent in the open arm by each animal expressing ChR2 or mCherry in the DRN during light-on and light-off conditions. In DRN-ChR2 mice, the duration spent in the open arm during light-on phase was significantly shorter than during the initial 3-minute light-off period (Friedman test with Wilcoxon signed-rank test for multiple comparison: $\chi^2$=11.556, *p*=0.003). **e** Schematics representation of the resident-intruder test. Blue light was intermittently activated and deactivated for 1-minute epochs over a duration of 10 minutes. **f** Duration of aggressive behaviors (including attacking and chasing; DRN-mCherry mice: Wilcoxon signed rank test; $Z_6$=-0.730, *p*=0.465; DRN-ChR2 mice: $Z_6$=-0.535, *p*=0.593) and non-aggressive behaviors (including grooming and sniffing; DRN-mCherry mice: $Z_6$=-1.05, *p*=0.917; DRN-ChR2 mice: $Z_6$=-1.153, *p*=0.249) during light-on and light-off conditions. **g** Schematics representation of the social memory test. The test mice were previously

habituated to one mouse (defined as the familiar mouse) for five minutes, and then a novel mouse was placed into the other chamber. For the test mice, blue light was intermittently turned on and turned off for 1-minute epochs over a duration of 5 minutes. **h** Both DRN-mCherry and DRN-ChR2 mice exhibited a significant preference towards the novel mice in the social memory test (Two-way repeated measures ANOVA with Bonferroni correction; $F_{1,10}$=6.942, *p*=0.025), with no significant difference observed between the two groups ($F_{1,10}$=0.000, *p*=1). **i** Schematics representation of the grip strength measurement. Each mouse underwent three blocks of testing, with two-hour intervals between blocks. In each block, mice were subjected to ten trials over approximately 6 minutes, divided into two equal 3-minute periods corresponding to light-on and light-off conditions. The maximum value recorded was used to define the mouse's grip strength.
**j** Photostimulation of DRN GABAergic neurons did not significantly alter grip strength in mice (Paired samples t-test; $t_8$=-0.894, *p*=0.397). **p* < 0.05, ***p* < 0.01. Data are expressed as mean ± SEM.

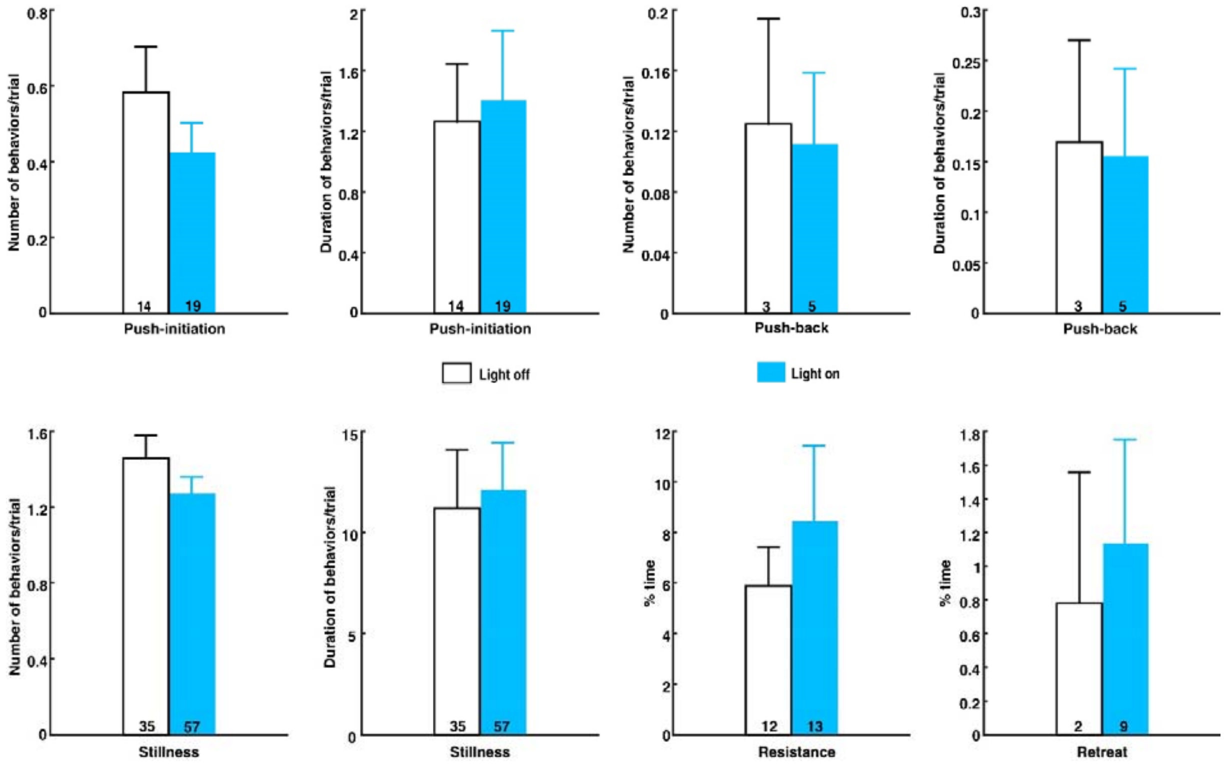


**Figure S3. The photostimulation protocol utilizing 473 nm blue light did not affect the behavior of DRN-mCherry mice in the tube test.** Optogenetic activation exhibited no significant impact on the number of push-initiations (Mann-Whitney U-test: U=461, *p*=0.253) or the duration of push-initiations (U=483, *p*=0.431). Similarly, there were no significant differences observed in the number of push-backs (U=532.5, *p*=0.865) or their duration (U=531, *p*=0.838), the number of stillness episodes (U=449, *p*=0.171) or their duration (U=405, *p*=0.089), as well as in the percentage of time spent resisting (U=430.5, *p*=0.080) and the percentage of time spent retreating (U=569.5, *p*=0.129). Data are presented as mean ± SEM.

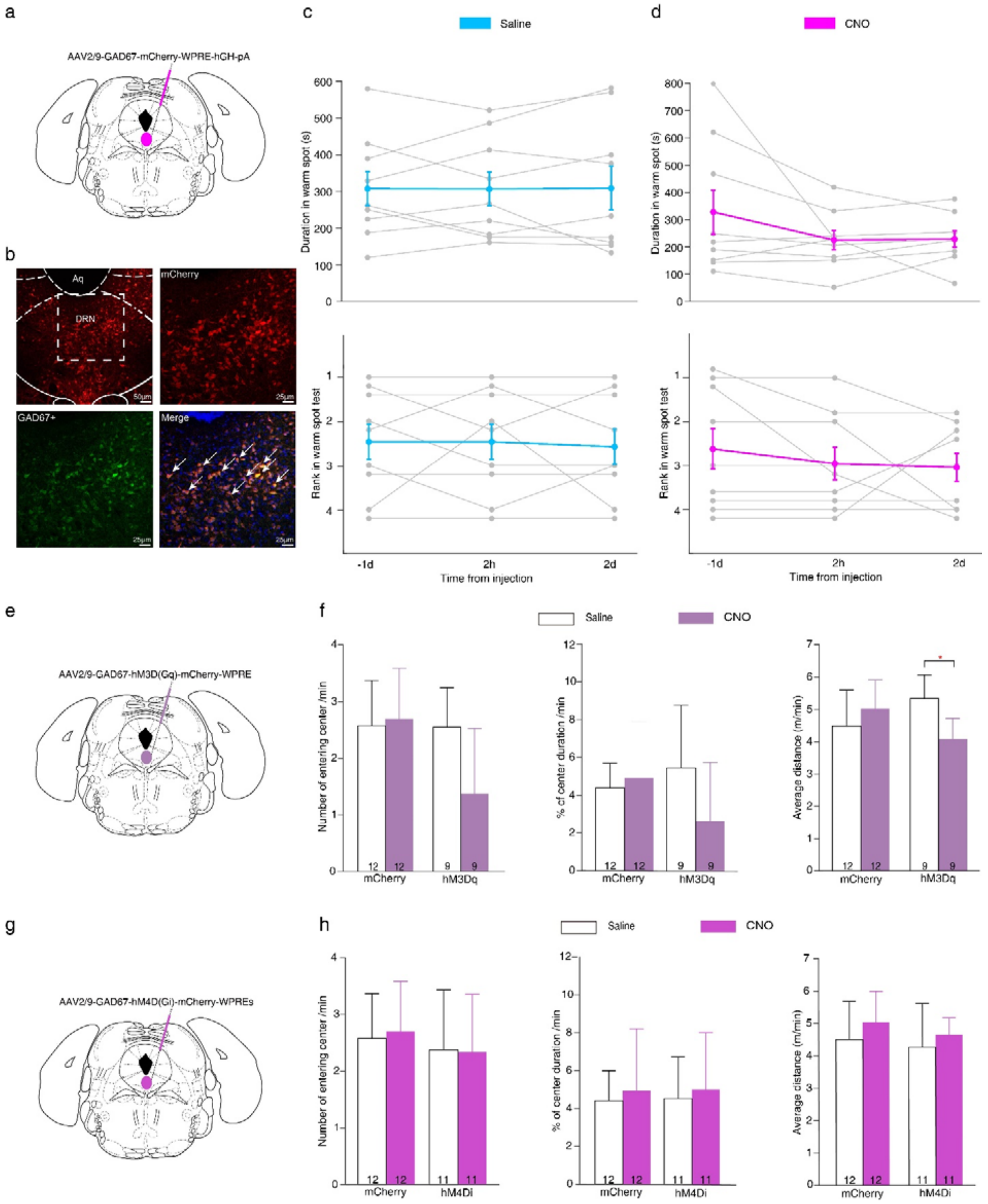

**Figure S4. Effects of chemogenetic manipulation on mouse behavior. a-d** The control virus did not significantly affect the performance of mice in the warm spot test. **a** Schematic representation of the injection site in the DRN for the control virus AAV2/9-GAD67-mCherry-WPRE-hGH-pA. **b** Representative photomicrographs of the DRN depicting mCherry (red, upper left) with the enlarged pictures for the boxed area showing mCherry expression (red, upper right), $GAD67^+$ (green, lower left), and their co-localization with DAPI (lower right). **c** Duration of warm spot occupancy (top; One-way repeated measures ANOVA with Bonferroni correction: $F_{2,16}$=0.004, *p*=0.996; n=9) and the rank in the warm spot test (bottom; Friedman test with Wilcoxon signed-rank test for multiple comparisons: $\chi^2$=0.4, *p*=0.819; n=9) in DRN-mCherry mice were assessed one day prior, 2 hours post, and 2 days post saline injection. Each gray folded line represents an individual animal, while the colored line indicates the average. Individuals of the same rank were arranged in parallel to facilitate the presentation of results. **d** Duration of warm spot occupancy (top; $F_{2,16}$=1.224, *p*=0.305; n=9) and rank in the warm spot test (bottom; $\chi^2$=1.368, *p*=0.504; n=9) in DRN-mCherry mice were assessed one day prior, 2 hours post, and 2 days post CNO injection. **e** Schematic representation of the injection site in the DRN for the virus AAV2/9-GAD67-hM3D(Gq)-mCherry-WPRE. **f** Chemogenetic activation significantly affected the mean distance traveled in the OFT (right; One-way repeated measures ANOVA with Bonferroni correction: $F_{1,8}$=7.646, *p*=0.024), but not the number of center entries (left; $F_{1,8}$=4.980, *p*=0.056) and the percentage of time spent in the center (middle; $F_{1,8}$=3.963, *p*=0.082). **g** Schematic representation of the injection site in the DRN for the virus AAV2/9-GAD67-hM4D(Gi)-mCherry-WPREs. **h** Following chemogenetic inhibition, the number of center entries (left; $F_{1,10}$= 0.045, p=0.837), the percentage of time spent in the center (middle; $F_{1,10}$=0.181, p=0.680), and the average distance traveled (right: $F_{1,10}$=1.133, p=0.312) in the OFT remained unaltered. $^*p < 0.05$. Data are expressed as mean ± SEM.

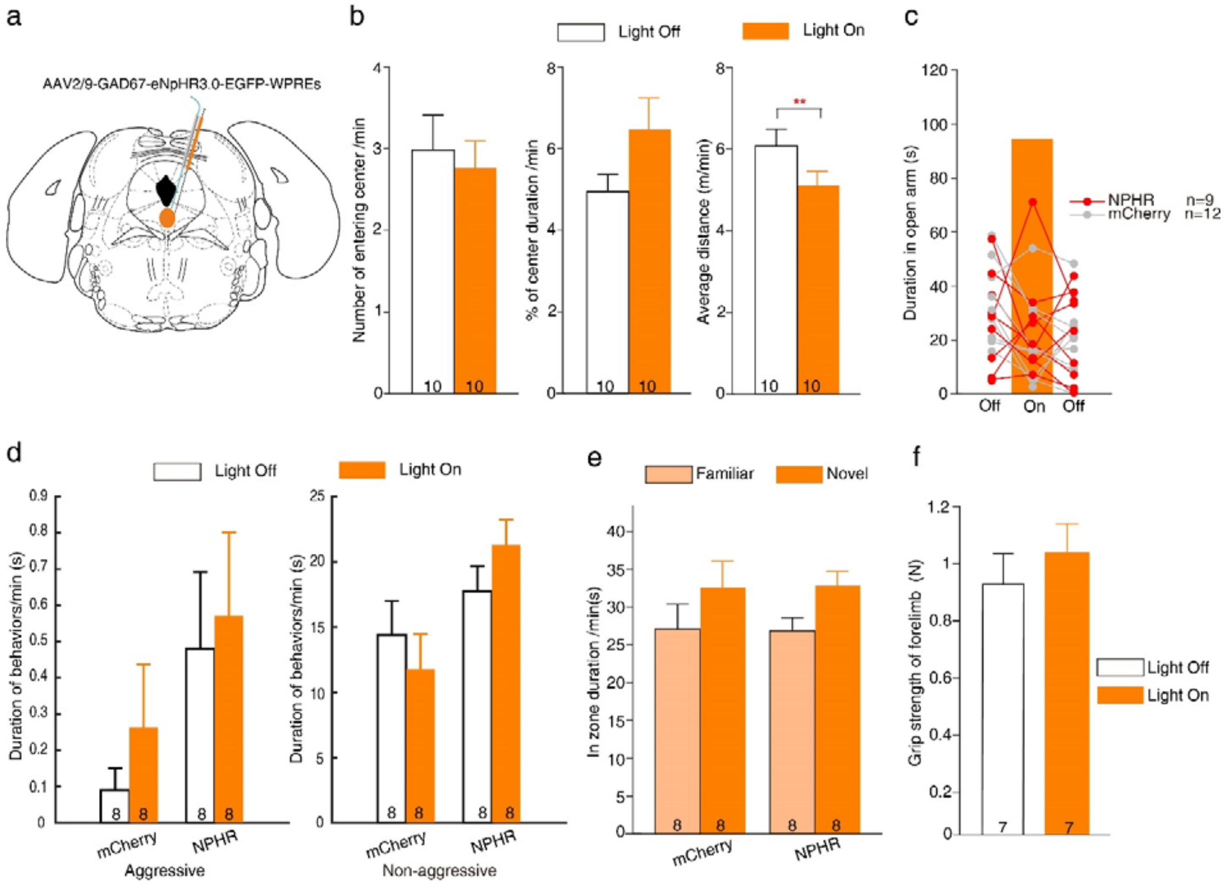


**Figure S5. Optogenetic inhibition of DRN GABAergic neurons did not significantly impact anxiety, aggression, social memory, or grip strength. a** Schematic representation of the injection site in the DRN for the virus AAV2/9-GAD67-eNpHR3.0-EGFP-WPREs. **b** Optogenetic inhibition of DRN GABAergic neurons did not affect the number of entries into the center (Paired samples t-test; left: $t_9$=0.823, *p*=0.432), the percentage of time spent in the center (middle; $t_9$=-2.113, *p*=0.064), but resulted in a significant reduction in average distance traveled (right: $t_9$=4.356, *p*=0.002). **c** Photostimulation did not alter the duration spent in the open arm by mice expressing either eNpHR or mCherry in DRN GABAergic neurons (Two-way repeated measures ANOVA with Bonferroni correction; for light condition: $F_{2,38}$=3.320, *p*=0.047, with no significant differences observed among the three light conditions; for group: $F_{1,19}$=0.345, *p*=0.564). **d** Both DRN-mCherry and DRN-NpHR mice showed normal levels of aggressive and non-aggressive behaviors towards novel mice in the resident-intruder test (Wilcoxon signed rank test; n=8 for each group; for DRN-mCherry mice: $Z_6$=-1.069 and *p*=0.285 for aggressive behavior, and $Z_6$=-1.400 and *p*=0.161 for non-aggressive behavior; for DRN-NpHR mice: $Z_6$=-0.314 and *p*=0.753 for aggressive behaviors, and $Z_6$=-1.820 and *p*=0.069 for non-aggressive behaviors), with no significant differences between the two groups (Mann-Whitney U test; for aggressive behaviors during light-on and light-off periods: U=27, *p*=0.584 and U=29, *p*=0.742; for non-aggressive behaviors during light-on and light-off periods: U=15, $p$=0.074 and U=30, $p$=0.834). **e** Both groups of mice exhibited a trend of preference towards novel mice in the social memory test (n=8 for each group, Two-way repeated measures ANOVA with Bonferroni correction; $F_{1,14}$=2.680, $p$=0.124), with no significant difference noted between the groups ($F_{1,14}$=0.000, $p$=1). **f** Photostimulation of DRN GABAergic neurons did not alter grip strength (Paired samples t-test: $t_6$=1.996, $p$=0.093). $^{**}p < 0.01$. Data are expressed as mean ± SEM.

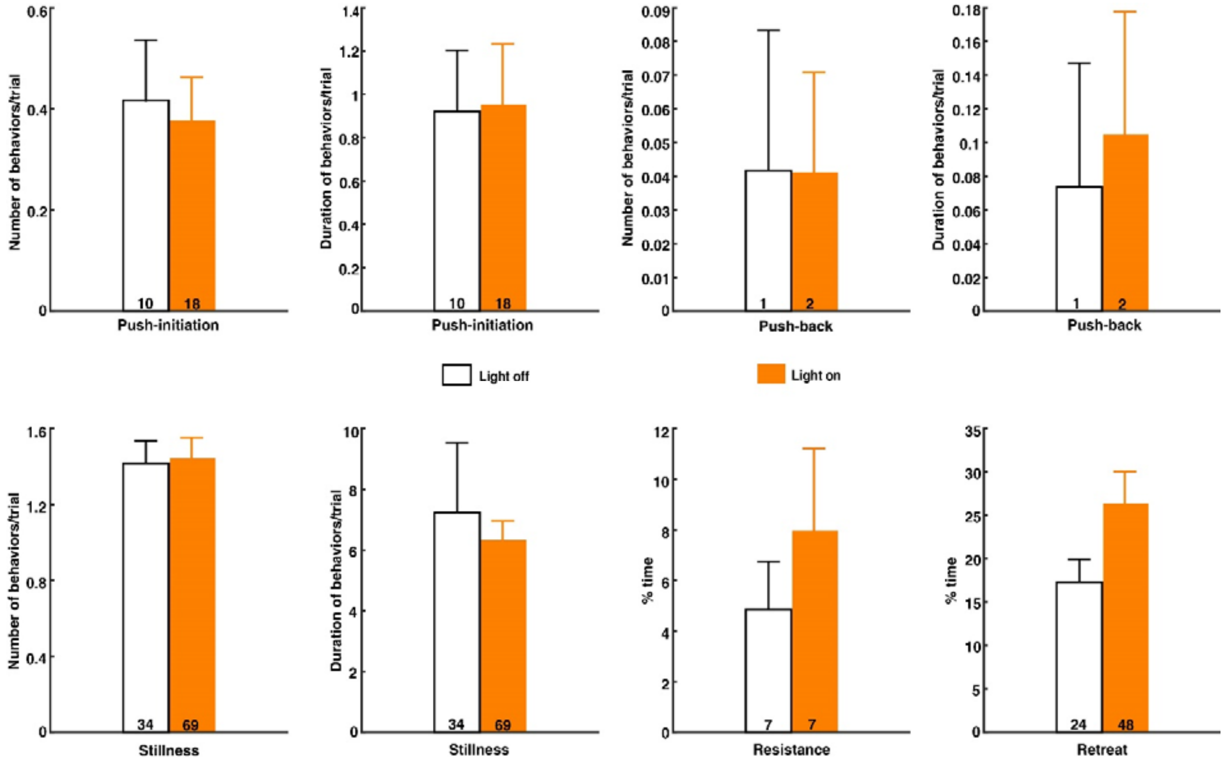


**Figure S6. The photostimulation protocol utilizing 589 nm orange light did not affect the behavior of DRN-mCherry mice in the tube test.** Optogenetic activation exhibited no significant impact on the number of push-initiations (Mann-Whitney U-test: U=546, $p$=0.664) or the duration of push-initiations (U=533.5, $p$=0.545). Similarly, there were no significant differences observed in the number of push-backs (U=576, $p$=1.000) or their duration (U=575, $p$=0.972), the number of stillness episodes (U=552.5, $p$=0.739) or their duration (U=434.5, $p$=0.091), as well as in the percentage of time spent resisting (U=504.5.5, $p$=0.216) and the percentage of time spent retreating (U=499, $p$=0.357). Data are presented as mean ± SEM.
